# Ex Vivo Expansion of Hematopoietic Stem and Progenitor Cells from Human Mobilized Peripheral Blood for Gene Therapy Applications

**DOI:** 10.64898/2026.04.08.716064

**Authors:** Erika Zonari, Matteo Maria Naldini, Matteo Barcella, Monica Volpin, Vincenti Francesca, Giacomo Desantis, Leila Hadadi, Carolina Caserta, Ilaria Galasso, Beatrice Martini, Francesca Tucci, Leonardo Ormoli, Ilaria Visigalli, Michela Vezzoli, Dejan Lazarevic, Ivan Merelli, Stephanie Z Xie, John E. Dick, Eugenio Montini, Bernhard Gentner

## Abstract

*Ex vivo* expansion of mobilized peripheral blood (mPB) hematopoietic stem cells (HSCs) represents a promising approach to advance cell and gene therapy strategies yet is hampered by loss of stem cell function when applying commonly used culture protocols. We performed in-depth characterization of mPB expansion cultures by single cell RNA sequencing, which highlighted differentiation trajectories with preservation of lineage fidelity in committed progenitors. Defining a putative HSC cluster allowed an estimation of transduction efficiency in *ex vivo* cultures, which correlated with long-term gene marking in xenografts and patients enrolled in a gene therapy study. We then developed a clinically translatable, GMP-compliant process to expand lentivirus (LV)-transduced HSCs from mPB of pediatric patients and adult donors, by biologically informed protocol improvements of cytokine supplementation, media choice, timing of LV transduction and combinations of small molecules preventing the activation of differentiation programs. Our optimized process outperforms validated state-of-the-art cord blood expansion protocols when applied to mPB. LV integration site analysis and genomic barcode-based clonal tracking provided definitive proof for symmetric HSC self-renewal divisions occurring during *ex vivo* culture. These results warrant clinical testing of this HSC transduction/expansion process in an upcoming clinical gene therapy trial for autosomal recessive osteopetrosis (EU CT 2024-518972-30).

**One Sentence Summary:** A mobilized peripheral blood HSC expansion protocol optimized for gene therapy allows robust polyclonal long-term engraftment of LV-transduced cells.

## INTRODUCTION

Continuous production of blood and immune cells throughout an individual’s life relies on a reserve of quiescent hematopoietic stem cells (HSC) that may dynamically respond to hematopoietic demand by undergoing self-renewal or multi-lineage differentiation. This triple ability is impressively recognizable after HSC transplantation, where a few thousand donor HSCs reconstruct and maintain a full blood and immune cell compartment in the recipient (*1, 2*). This functional potential of HSCs is intimately linked to a complex bone marrow (BM) niche (*3*). Transplantation of unmodified or genetically engineered HSCs can be an effective treatment for hematological malignancies and non-malignant disorders (*4*) but may be limited by availability of HLA-matched HSCs (in allogeneic settings) and yield of successfully genetically engineered cell products. *Ex vivo* expansion of HSCs aims to overcome such limitations by increasing numbers of functional HSCs/progenitors and/or enhancing their engraftment potential *in vivo*, thus potentially widening access to HSC-based treatments.

One clinically relevant application for *ex vivo* expansion is allogeneic umbilical cord blood (CB) transplantation, where the small size of most CB units precludes their safe clinical use, especially in adults. This bottleneck could be overcome by *ex vivo* expansion before infusion into patients, allowing for a potentially broader choice of CB units with improved HLA matching (*5–8*). Successful approaches have leveraged cytokine stimulation to promote CB HSC proliferation and survival, combined with molecules that counteract HSC commitment to differentiation, the default program triggered under these culture conditions, during which protein synthesis, signaling and stress responses are upregulated (*9–11*). Molecules shown to help maintain CB HSC modulate histone modifiers (*12–16*), the aryl hydrocarbon receptor pathway (*17, 18*), metabolism and organelle biology (*19–21*), and Notch signaling (*5, 22*). Many of these strategies have shown functional evidence for some degree of HSC expansion when using CB as a source. Still, it remains largely unclear whether the combination of molecules acting through different mechanisms could further improve HSC expansion, and whether these protocols perform similarly on other HSC sources. Another approach has focused on chemically defined culture media titrating the activation of essential signaling pathways and minimizing batch variability and contaminants often associated with products of biological origin (*23*). Experimental research approaches are also exploring more complex systems, such as 3D cultures in hydrogels (*24*) or co-culture with HSC-supporting scaffolds or stromal cells mimicking the bone marrow niche (*25, 26*). While promising, these models present significant limitations in terms of translational potential towards clinical applications, which require scalability and compliance with good manufacturing practice (GMP) standards.

Another emerging application for *ex vivo* expansion is gene therapy (GT) of the patient’s own HSC, which is associated with less morbidity and potentially higher efficacy compared to allogeneic transplantation, when it comes to the treatment of select monogenic disorders (*27*). In GT protocols, HSCs are genetically modified by vector mediated addition of a therapeutic gene or edited for correction of genetic defects (*28*). However, GT protocols may reach suboptimal gene transfer/correction efficiencies or may involve manipulation of autologous patient HSCs with low yield (e.g. in very young pediatric patients ineligible for leukapheresis or individuals with poor stem cell mobilization potential), or disorders where HSC quantity and quality are inherently compromised, such as reversible niche defects, bone marrow failure syndromes or chronic inflammatory conditions. *Ex vivo* expansion applied to GT protocols may allow to expand and/or select for genetically modified HSCs to achieve sufficient transgene-carrying cells for therapeutic efficacy. Furthermore, expansion of corrected HSCs may allow downscaling of transduction and editing steps reducing manufacturing costs. Hence, there are multiple arguments for applying *ex vivo* HSC expansion protocols in the autologous HSC gene therapy context. Unfortunately, autologous CB units are generally not available as an HSC source. Although a previous study has examined whether HSC expansion protocols developed for CB may sustain stemness in mobilized peripheral blood (mPB) HSCs (*29*), the preferred source for gene therapy, functional evidence for *ex vivo* expansion of mPB or BM HSC remains limited. Likewise, the impact of the genetic engineering step on *ex vivo* expansion protocols is poorly characterized.

Here, we performed an in-depth characterization of lentiviral vector (LV)-transduced, *ex vivo*-expanded mPB-HSC from pediatric patients enrolled in a gene therapy trial (NCT03488394) and from adult healthy donors, including single cell RNA sequencing (scRNAseq) and clonal graft analysis in primary and secondary xenografts. While showing robust evidence for symmetric HSC division in *ex vivo* culture, we found that net HSC expansion is considerably lower compared to the data described for CB. To improve net mPB HSC expansion, we explored combinations of HSC maintaining molecules and undertook a hypothesis-driven, iterative optimization of the culture protocol towards the development of a clinically applicable process to produce LV-transduced, *ex vivo* expanded HSCs from mPB.

## RESULTS

### Net HSC maintenance in UM171-expanded mPB CD34+ cells from healthy donors and gene therapy patients

To test previously described *ex vivo* expansion protocols, developed for umbilical CB (*12*) on clinically relevant HSC sources in a gene therapy context, we cultured hematopoietic stem and progenitor cells (HSPC) from n=5 mPB drug products (DPs) from a pediatric gene therapy trial (NCT03488394) for mucopolysaccharidosis type I (MPS) (*30*) for 7 to 8 days using commercial media supplemented with early acting cytokines (SFT6: Stem cell factor, 100ng/mL; FLT3 ligand, 100ng/mL; Thrombopoietin, 50ng/mL; Interleukin 6, 50ng/mL), with or without the epigenetic modifier UM171 (Fig.1A). DP-derived cells expanded 8- to 27- fold (Fig.1B) and, as expected, the addition of UM171 increased the proportion of cells with a primitive immunophenotype compared to cytokine-only cultures (Fig.1C).

**Figure 1.**
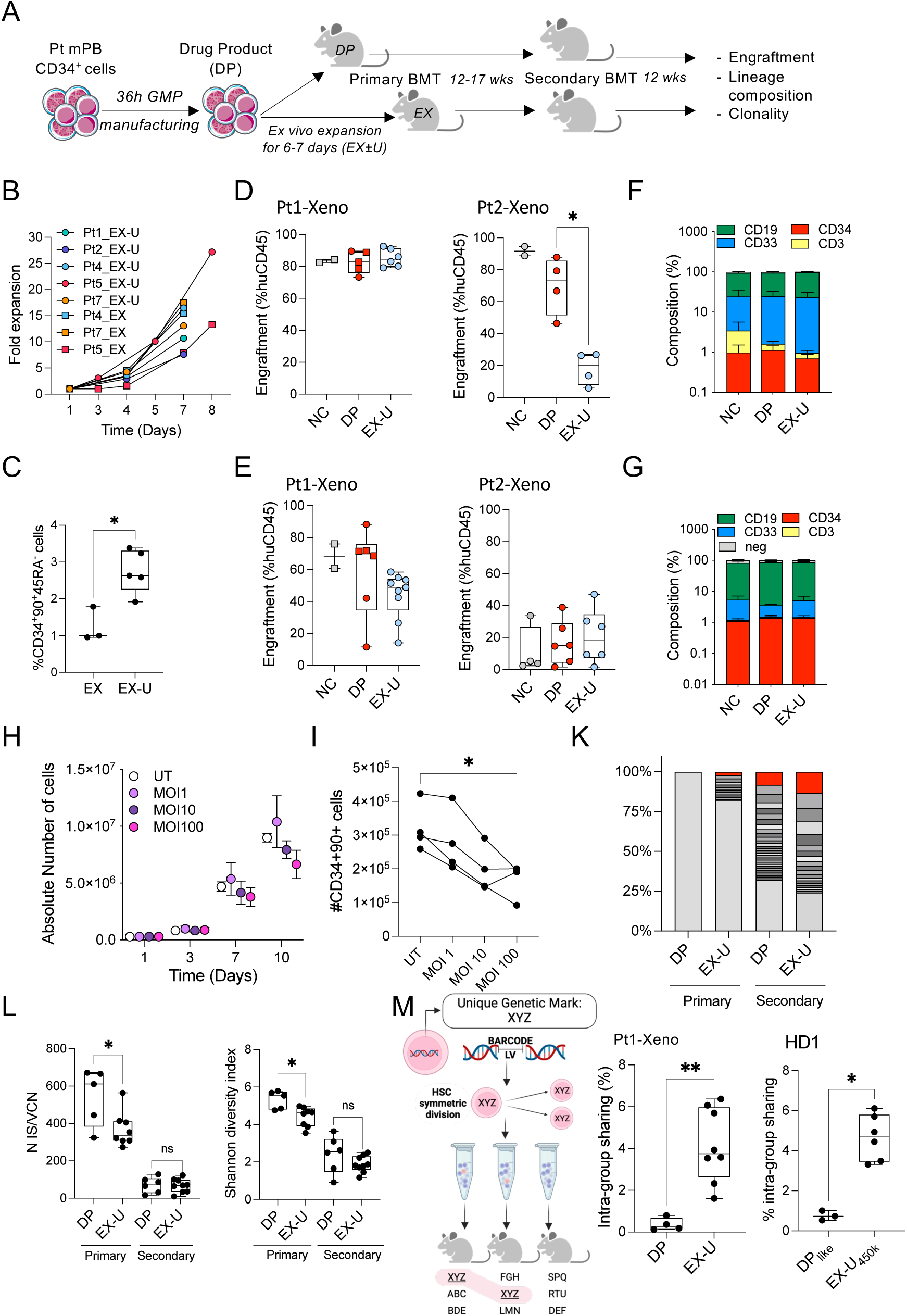
Net HSC maintenance in UM171-expanded mPB CD34+ cells. (**A**) Schematic representation of the experimental design. (**B**) *Ex vivo* expansion of drug products (DP) from different MPS1 patients (Pt) for 6-7 additional days in the presence (dot; EX-U) or absence (square; EX) of UM171 (35nM). Fold expansion is calculated as a ratio between final cell count on day 7/8 with respect to seeding on day 1. (**C**) Fraction of HSC-enriched CD34+CD90+CD45RA- cells at the end of expansion culture of the samples from panel (B). *:p<0.05, Mann-Whitney test. (**D**) Human CD45⁺ cell engraftment in the bone marrow of NSG mice at 12 weeks post i.v. injection of the following mPB-derived HSPC products from 2 MPS1 patients: Pt1-Xeno (i) 400,000 non-cultured CD34⁺ cells (NC; *n* = 2), (ii) 400,000 DP cells (DP; *n* = 5), or (iii) 4,000,000 *ex vivo* expanded cells (outgrowth of 400,000 DP cells; EX-U; *n* = 6). Pt2-Xeno (i) 150,000 non-cultured CD34⁺ cells (NC; *n* = 2), (ii) 150,000 DP cells (DP; *n* = 4), or (iii) 1,100,000 *ex vivo* expanded cells (outgrowth of 150,000 DP cells; EX-U; *n* = 4). *:p<0.05, Mann-Whitney test. BM cells were obtained as terminal procedure or BM aspirate for the mice injected with cells from Pt1 and Pt2, respectively. See Fig.S3 for the endpoint analysis of Pt2-Xeno. (**E)** Secondary transplantation of 2 and 1 × 10⁶ CD34^+^ cells isolated from primary NSG recipients engrafted with HSPC from Pt1-Xeno and Pt2-Xeno, respectively. BM engraftment of secondary recipients was assessed at 12 weeks post transplantation, respectively. Each dot represents an individual mouse **(F, G**) Lineage composition of the human CD45^+^ cell graft from panels D, E, respectively. B cells: CD19+; myeloid cells: CD33^+^; T cells: CD3^+^; CD34^+^: HSPC cells; lineage negative: CD19^-^CD33^-^CD3^-^CD34^-^. Shown is the mean ± SEM from the pooled individual mice from Pt1-Xeno and Pt2-Xeno. (**H**) *Ex vivo* expansion (mean ± SEM) of mPB CD34^+^ cells from n=4 adult donors under EX-U conditions following transduction with a lentiviral vector at increasing multiplicities of infection (MOI). UT, non-transduced cells. (**I**) Absolute number of CD34^+^90^+^ cells on day 7 of expansion culture from (H). Each line represents a donor (*P<0.05, Kruskal-Wallis test with Dunn’s multiple comparison). (**K**) Relative abundance of unique lentiviral insertion sites (LVIS) from DP and EX-U groups from Pt1-Xeno in primary and secondary recipient mice. Bottom light grey stacks indicate the sum of insertions with <1% relative abundance, middle stacks indicate insertions >1% abundance, where the top abundant insertions are highlighted in red. (**L**) Left panel: box plots indicating number of insertion sites normalized on vector copy number. Right panel: box plots indicating the Shannon diversity index as measure of clonal population diversity. Each dot indicates an individual mouse of the treatment group: DP and EX-U in primary and secondary recipient respectively (*P<0.05, Mann-Whitney test). (**M**) Left panel: Schematic representation exploiting unique LVIS or lentiviral barcodes (LVBC) to prove symmetric HSC division in culture. Right panel: Boxplots showing the proportion of shared LVIS (Pt1-Xeno) or shared LVBCs (HD1) between at least 2 individual mice receiving the same d0 equivalent CD34+ cell doses from the indicated HSPC products (intra-group sharing). Each dot represents the BM huCD45+ graft of an individual mouse (primary transplant) at the experimental endpoint. (*P<0.05, **P<0.01, Mann-Whitney test). Further details on HD1 are presented in Supplemental Fig.1. In all box plots shown in the figure, the whiskers represent the minimum and maximum values of the data.

To assess the repopulating potential of MPSI-H patient-derived DPs expanded in the presence of UM171 (EX-U), we measured human cell engraftment and lineage composition at 12 weeks after xenotransplantation into primary (Fig.1D,F) and secondary recipient immunodeficient mice (Fig.1E,G). EX-U cells were compared, at matched pre-expansion cell doses (“day 0 equivalent”), to their respective non-expanded DPs and the non-cultured (NC) CD34+ cells used as starting material for the *ex vivo* engineering/culture process. All conditions resulted in detectable human cell engraftment in the bone marrow of primary and secondary recipients (Fig.1D,E), with multilineage output (Fig.1F,G). *Ex vivo* culture had a different impact on short-term (ST-) HSC function from the DPs of the 2 patients (Pt), resulting in a similar-sized primary graft for the DP from Pt1 (Fig.1D, left), but significantly reduced engraftment for the DP from Pt2 (Fig.1D, right). Nevertheless, proficient transfer of the grafts from the 2 patients to secondary recipients (Fig.1E,G) suggested that *ex vivo* expanded cells maintained long-term (LT-) HSC potential.

We hypothesized that the reduced primary engraftment of EX-U cells from Pt2 in xenotransplantation assays could be related to lentiviral (LV) transduction when coupled to *ex vivo* expansion, as this patient had the most highly transduced DP in the study, with an average vector copy number per cell (VCN) of 5.2, compared to a median VCN of 2.2 when considering all the 8 treated patients (*30*). To further test this hypothesis, we transduced mPB CD34+ cells from 4 healthy adult donors with increasing LV doses observing, similarly to the DP of Pt2, a dose-dependent decrease in cell growth during *ex vivo* culture (Fig.1H), with concomitant reduction in the absolute number of immunophenotypically more primitive cells (Fig.1I). Thus, *ex vivo* culture may amplify well-described HSPC stress responses induced by genetic engineering technologies particularly impacting ST-HSC function (*31*), investigated in more detail below.

Next, we analyzed lentiviral vector integration sites (LVIS) to determine the clonal structure of EX-U versus DP-derived grafts from Pt1. LVIS distribute semi-randomly across the genome. Hence, each individual transduced clone is characterized by one or more unique integration site(s) stably transmitted to the progeny. Xenografts derived from DP and EX-U had a highly polyclonal pattern in the primary mice, without detection of dominant clones (Fig.1K). As expected from a procedure enriching for long term (LT)-HSC, secondary grafts had a less polyclonal composition in both groups but still showed no signs of clonal dominance (Fig.1K). Quantitative comparison between EX-U and DP showed that primary, but not secondary grafts originating from EX-U had significantly fewer unique integration sites and a lower Shannon diversity index, suggesting some loss of ST-HSC activity also for cells from Pt1 during *ex vivo* expansion, but preserved long-term engraftment potential (Fig.1L).

To confirm the broader applicability of our expansion protocol to adult mPB, we compared the engraftment potential of EX-U and DP-like HSPCs from an adult healthy donor (HD1) after transduction with a highly diverse LV barcode library, simulating a gene therapy context and allowing simultaneous clonal graft analysis by an independent methodology (Fig.S1A). EX-U were xenotransplanted at decreasing day 0-equivalent cell doses, resulting in a dose-dependent decrease in peripheral blood human cell engraftment (Fig.S1B). After 16 weeks from xenotransplantation, EX-U showed comparable engraftment to DP-like cells when injecting the same day 0 dose equivalent (Fig.S1C), yet lower but still detectable multi-lineage engraftment (Fig.S1D) when decreasing the dose to 33% and 9% of the DP-like input dose. We assessed graft clonality by 2 methods, either by LVIS or by quantifying the abundance of individual barcodes introduced into the cells by the lentiviral vector barcode (LVBC) library. Both methods consistently showed polyclonal reconstitution even at the intermediate expanded cell dose, with a concordant and strikingly similar clonal abundance pattern supporting the robustness of the 2 methods (Fig.S1E,F; Table S1). Clonal diversity and the estimated number of human repopulating clones per mouse was equivalent or higher in the EX-U compared to the DP-like group at equivalent day 0 starting cell doses in line with at least LT-HSC maintenance during *ex vivo* culture (Fig.S1G,H). To prove that some of these HSC had undergone symmetric self-renewal divisions during *ex vivo* expansion, we identified unique LVIS or LVBCs from Pt1 and HD1, respectively, that could be detected in different mice injected with the same cell therapy product (Fig.1M). Only background levels of such shared clones were detected in mice transplanted with DP(-like) cells, where cell counts remained stable during the short duration of culture, while 2-7% of clonal markers were shared between mice that received EX-U cells (Fig.1M; Table S1).

Overall, these data suggest that the UM171-culture allowed net maintenance of adult and pediatric HSC from mPB, likely due to some HSCs undergoing symmetric division *ex vivo* and others being lost during the culture process. *Ex vivo* culture did not result in clonal dominance, preserving clonal diversity.

### *Ex vivo* culture of mPB CD34+ cells mostly expands committed progenitors with preserved lineage fate and allows prediction of in vivo DP characteristics by scRNAseq

To characterize mPB CD34+ cell population dynamics during EX-U culture, we performed single cell RNA sequencing (scRNAseq) on cells from 2 adult healthy donors. CD34+ cells from donor 1 were analyzed before culture, on day 4 and on day 8 of expansion. For donor 2, in addition to the CD34+ bulk expansion culture, part of the CD34+ cells were FACS-sorted into subpopulations enriched in multipotent HSPC (CD34+CD38-), committed granulocyte-monocyte progenitors (GMP) or megakaryocyte-erythrocyte progenitors (MEP), and each subpopulation was marked by transduction with a LV expressing a different fluorescent protein for tracing back the population of origin prior to remixing the subpopulations together for expansion (Fig.2A). Colony-forming cell assay confirmed functional enrichment of GMP and MEP post sorting (Fig.2B). A single object containing single cell transcriptomes from all samples was generated following batch correction and dimensionality reduction. Unsupervised clustering was performed, and cellular states were manually annotated (Fig. 2C; Table S2). Compared to non-cultured cells, presumably more enriched in primitive cells, HSC and multi-lymphoid progenitors (MLP) progressively diminished over time in culture, while progenitor cell states, erythroid/megakaryocytic, actively cycling myeloid progenitors and mast cell precursors (MCP) increased (Fig.2D). These data suggest that expansion culture mainly supported the growth of mPB-derived progenitors, rather than more primitive HSC, which are progressively diluted during culture. We next asked whether committed progenitors were able to maintain lineage fate during culture, or whether there was plasticity. The expansion culture output of day 0 sorted GMP and MEP faithfully mapped on the myeloid/monocytic and erythroid/megakaryocyte progenitor clusters, respectively (Fig.2E,F). The day 0 CD34+CD38-progeny lacked the most differentiated states, showed a bias for the myeloid progenitor area and maintained a sizeable fraction of cells within the HSC cluster (Fig.2F), in line with our previous data advocating the CD34+CD38- population as an HSC-enriched starting population for expansion cultures (*29*). Taken together, these data support that HSPC fate is maintained during expansion culture.

**Figure 2:**
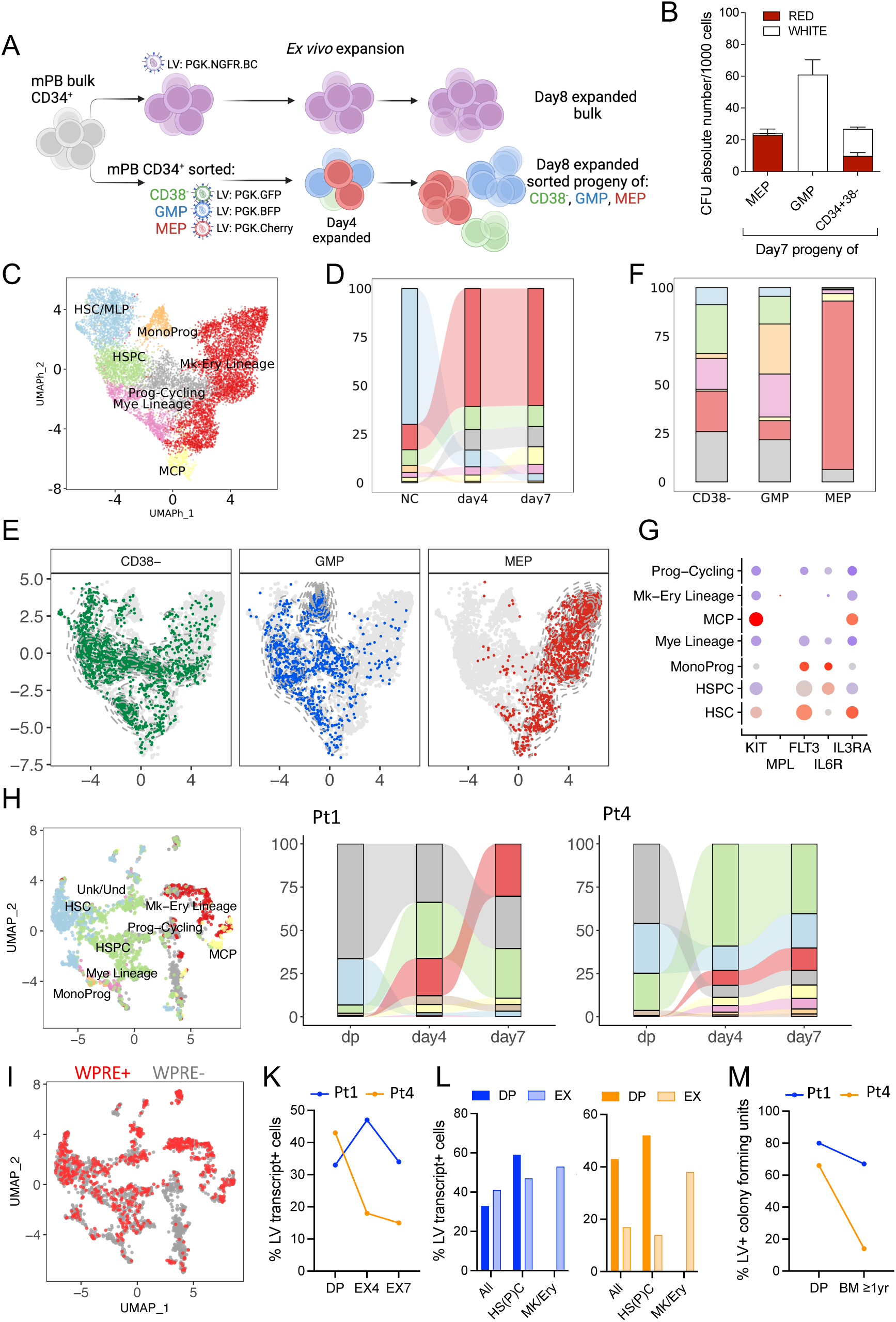
Characterization of expansion cultures by scRNAseq. **(A)** Adult mPB CD34+ cells from n=2 healthy donors were expanded with UM171 in bulk following transduction with a barcoded LV library (upper panel) or after marking FACS-sorted GMP (CD34+CD38+CD45RA+ or CD34+CD38+CD45RA-BAH1-CD71-), MEP (CD34+CD38+CD45RA-CD71+) or multipotent HSPC (CD34+CD38-) with a color-coded LV expressing a blue (BFP), red (Cherry) or green (GFP) fluorescent protein, respectively (lower panel). Cells were harvested on day 4 and/or day 8 for scRNAseq library preparation. **(B)** Colony-forming cell assay on sorted progeny of GMP, MEP and CD38- cells after 7 days of expansion. Absolute number of colonies forming units (CFU) with red (BFU-E) or white (CFU-G/M) morphology are shown for the 3 populations of origin (mean ± SEM, n=5 technical replicates). **(C)** Uniform Manifold Approximation and Projection (UMAP) encompassing the samples described in (A) with manually curated cell type annotation, as described in methods. HSC/MLP, hematopoietic stem cells/multi-lymphoid progenitors; HSPC, hematopoietic stem/progenitor cells; Mye, myeloid lineage progenitors; Mono, monocyte progenitors; MCP, mast cell progenitors; Mk-Ery, megakaryocyte/erythroid lineage progenitors; Prog-Cycling, progenitors predicted to be in active phases of the cell cycle. **(D)** Alluvial plot showing population frequencies over time, calculated within each timepoint. NC (non-cultured sample – day0). **(E)** Distribution of the expanded Day 7 progeny of FACS-sorted cell populations across UMAP embedding according to cell population of origin and **(F)** relative frequency of scRNAseq-based cell type annotations as defined in panel (C). (**G)** Expression of key cytokine receptors in the indicated cell type annotations. The color palette (from blue to red) represents the z-score of receptor expression (from low to high, respectively), while the size of the dots represents the percentage of cells expressing the indicated cytokine receptor transcript in the indicated population. (**H)** Pediatric mPB DPs from n=2 MPS1 patients (Pt1, Pt4) were sorted for CD34+CD90+CD45RA- cells (dp = day 1) or expanded for 4 (EX4) and 7 days (EX7) in UM171 before sorting into the CD34+CD90+CD45RA- cell fractions. scRNAseq was performed on the HSC-enriched subpopulations at the 3 timepoints. The UMAP on the left shows automated cell type annotation using scGate leveraging the adult mPB dataset described in panels (A-G) as reference (see methods for details). The alluvial plots on the right show time-dependent changes in population frequencies. Note that CD34+CD90+CD45RA- cells, an HSC-enriched subset of the cultures, was analyzed here. (**I)** Cells with lentiviral (LV)-derived transcripts (*WPRE+*) are projected on the UMAP plot, as a surrogate for estimating transduction efficiency. (**K)** Proportion of LV transcript+ cells over time in the 2 patients. Transduction efficiency overtime and across major populations. (**L**) Proportion of LV transcript+ cells in the DP (intensely colored bar) vs. the expanded cells (faintly colored bars; EX4 and EX7 samples were aggregated) for the entire population (All) or within the indicated, scRNAseq-defined subpopulations. HSC and HSPC were aggregated to obtain a representative number of cells for all conditions. Left graph (blue bars), Pt1; right graph, orange, Pt4. (**M)** Transduction efficiency assessed by digital droplet PCR on individually plucked colony forming units (gold standard assay), either on the DP (bulk) or in the BM of the patients at 1 year after gene therapy, as reported in Gentner et al. (*30*).

To evaluate the overall impact of *ex vivo* culture on HSPC transcriptomes, we identified longitudinally deregulated genes common across HSPC populations in the single-cell RNA sequencing dataset and assessed for enrichment of hallmark gene expression signatures (Fig.S2; Table S2). Hallmarks enriched from upregulated genes overtime included those related to proliferation, oxidative metabolism, mTORC1 signaling, and lipid metabolism, while hallmarks associated with inflammation and stress responses were broadly suppressed. To evaluate the influence of cytokine supplementation on culture dynamics, we assessed the expression of key cytokine receptors on different cell populations in scRNAseq. In addition to KIT and FLT3, also IL3RA was highly expressed in the most primitive annotated HSC subpopulation, while IL6 receptor was predominantly detected in the HSPC and monocyte progenitors (Fig.2G). These findings prompt additional investigations on the role of IL6 and IL3 supplementation during mPB expansion (see below).

Next, we hypothesized that in-depth characterization of expansion cultures by single-cell transcriptomics could aid in understanding DP characteristics where current standard release assays may not fully correlate with *in vivo* behavior. Although HSC-GT treatment resulted in encouraging metabolic correction and early clinical outcomes in all treated patients as of an interim analysis in all patients, the *in vivo* VCN stabilized in 3 out of 8 patients from the MPS trial at values below 0.3, whilst in vitro assays on the drug product measured a transduction efficiency >60%, and VCN ≥1 (*30*). Xenotransplantation accurately recapitulated steady-state VCN in patients (Fig.S2B) thus confirming that the VCN drop upon transplantation in the 3 patients was not due to reduced conditioning. We performed scRNAseq on CD34^+^CD90^+^CD45RA^-^, HSC-enriched populations from DP, day 4 EX-U and day 7 EX-U cultures from two representative MPS patients, namely Pt1 (correspondence between in vitro/in vivo VCN) and Pt4 (VCN drop upon transplantation). Cell populations were annotated by label transfer from the dataset of Fig.2B (Fig.2H; Table S3). The DPs from the 2 patients had similar compositions and EX-U showed the expected population dynamics, with a contraction of the HSC population (more in Pt1 than in Pt4) and an expansion of HSPC and Ery/Mk progenitors (more in Pt1). By assessing expression of LV-specific transcripts (*WPRE)* at single cell resolution (Fig.2I) we were able to estimate the percent of LV+ transcriptomes over time and within specific cell clusters as a putative surrogate for transduction efficacy (Fig.2K; Table S3). Strikingly, we recapitulated a drop in WPRE+ cells during EX-U for Pt4, but not Pt1, and this was evident in HS(P)C but not Ery/MK progenitors (Fig.2L; Table S3), resembling the drop in LV+ CFU in the BM of the patient with respect to the DP (Fig.2M). These data highlight the potential of scRNAseq of expansion cultures to predict upfront critical DP characteristics such as gene transfer into LT-HSCs.

### Iterative optimization of mPB CD34 cell expansion for gene therapy applications

To further optimize the EX-U protocol for gene therapy applications, we took an iterative approach. First, we asked whether IL6 could be substituted with IL3 based on receptor expression in cultured HSC (see Fig.2G). Culture in the presence of IL3 increased expansion by >3 fold (n=3 volunteer mPB donors), while reducing the proportion of CD34+ cells, leading, nevertheless, to a net gain in the absolute number of CD34+ cells (Fig.3A,B). On the other hand, IL6 tended to increase the absolute number of more primitive CD34+CD90+ cells (Fig.3C), and this effect was maintained when combined with intermediate doses of IL3 (condition G, SFT63 combo). Xenotransplantation confirmed a trend for improved engraftment upon EX-U with SFT63, suggesting that intermediate doses of both IL6 and IL3 are beneficial (Fig.3D).

**Figure 3.**
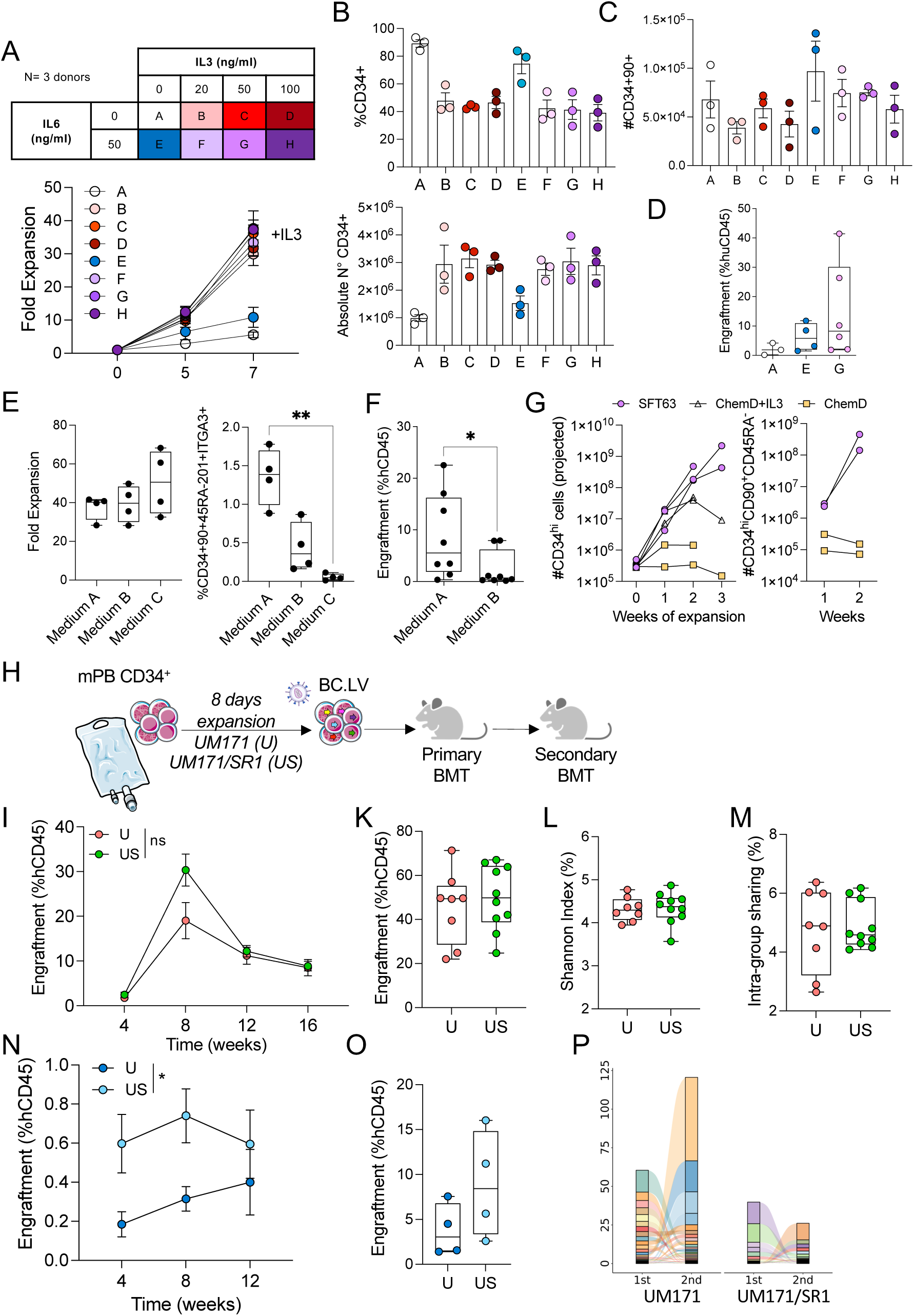
Expansion protocol optimization for mPB HSPC gene therapy applications. (**A**) (Top) The matrix outlines the IL-6 and IL-3 doses (ng/mL) used in the different culture combinations (identify by a different letter A-H), tested on three healthy donors. All conditions contained SCF 100ng/ml; Flt3L 100ng/ml; TPO 50ng/ml. (Bottom) Fold expansion over time (mean ± SEM) from three different donors cultured with the different IL-3 and IL-6 concentrations. (**B**) The percentage (top) and absolute number (bottom) of CD34^+^ cells measured by flow cytometry after 7 days of expansion under different conditions. Bars represent the mean ± SEM of individual donor values, shown as dots. (**C**) Absolute number of CD34^+^CD90^+^ cells measured by flow cytometry after 7 days of expansion under different conditions. Bars represent the mean ± SEM of individual donor values, shown as dots. (**D**) Percentage of human CD45^+^ cells present in the bone marrow (BM) of mice transplanted (n=3-6) with cells derived from the indicated culture conditions, 17 weeks post-transplant. The mice were transplanted using a pooled sample from the three donors for each condition, each mouse receiving a limiting dose of 35,000-39,000 day0 equivalent cells (Kruskal-Wallis test). (**E**) Different commercial media were compared (A: SCGM Cell Genix; B: StemSpan StemCell Technologies; C: StemPro GIBCO). Boxplot (left) showing the in vitro fold expansion measured after 7 days of culture of mPB CD34+ cells from four different healthy donors. In all tested conditions, the culture media were supplemented with the same cytokine cocktail (100 ng/mL SCF, 100 ng/mL Flt3L, 50 ng/mL TPO, 50 ng/mL IL-6, 50 ng/mL IL-3). The percentage of primitive CD34^+^CD90^+^CD45RA^-^CD201^+^ITGA3^+^ cells (right) was measured by flow cytometry after 7 days of expansion in different media. Each dot represents an individual donor (Kruskal-Wallis test with Dunn’s multiple comparison, *p<0.05, **p<0.01). (**F**) Boxplot showing the percentage of human CD45+ cells engraftment in the bone marrow (BM) of mice transplanted (n=8) with cells cultured in medium A or medium B at 16 weeks post-transplant. Each dot represents a single mouse, mice were transplanted using a pool from different donors for each condition, with 48,000-57,000 day0 equivalent cells injected per mouse (Mann-Whitney test, *P<0.05). (**G**) CD34^+^ cells from n=3 adult mPB donors were expanded over 3 weeks in medium A supplemented with SFT63 cytokines, UM729 and SR1 (SFT63), or a chemically defined medium (ChemD, see Methods), with or without IL3. The projected absolute number of CD34^high^ cells (left) and more primitive CD34^hi^CD90^+^CD45RA^-^ cells (right) over a 3-week culture period is shown, taking into consideration the proportion of cells that has been removed during consecutive passages. Connected dots refer to longitudinal data from an individual donor and the indicated condition. (**H**) Schematic representation of the experimental design. Different compound combinations were tested: UM171 (35 nM) alone (U) or in combination with SR1 (US) (750 nM). In both conditions, cells were cultured in SGCM with the optimized cytokine cocktail described above. (**I**) Percentage of human CD45^+^ cells (mean ± SEM) present in the peripheral blood (PB) of U (n=8) and US (n=10) conditions at the indicated number of weeks post-transplant. ns, not significant (U vs US, mixed effects analysis). (**K**) Percentage of human CD45^+^ cells present in the bone marrow (BM) at 17 weeks post-transplant. (**L**) Box plots indicating the Shannon diversity index as measure of clonal population diversity. Each dot indicates an individual mouse of the U and US primary recipient groups. (**M**) Sharing of barcodes within the same treatment group, measured as the percentage of shared barcodes from individual mice (dots) of the same treatment group. (**N**) Percentage of human CD45^+^ cells (mean ± SEM) present in the peripheral blood (PB) of U and US (n=4) conditions at the indicated number of weeks post-secondary transplant. The mice were transplanted using a pooled sample from bone marrow of primary recipient depleted of murine cells. The amount of CD34+ cells transplanted into secondary recipients ranged between 1-1.3×10^6^ CD34^+^ cells. *, p<0.05 (U vs US, Two-way Anova). (**O**) Boxplot with percentage of human CD45^+^ cells present in the bone marrow (BM) of mice transplanted (n=4) with cells derived from a primary recipient 17 weeks post-transplant. (**P**) Alluvial plots showing the frequency of shared barcodes between primary and secondary transplants. The barcodes are arranged from top to bottom by decreasing frequency in the primary transplant, considering only those that are also found in the secondary graft. The frequency of each barcode is calculated as the mean frequency in mice bearing the barcode. Left and right alluvial plots show the sharing in the UM171 (U) and UM171/SR1 (US) conditions, respectively.

Second, EX-U was tested with SFT63 on mPB CD34+ cells from n=4 volunteer donors in the context of different commercial media. While fold expansion was similar, medium A (SCGM, CellGenix) best maintained cells with a primitive immunophenotype in culture, which translated into significantly higher engraftment as compared to medium B (Fig.3E,F). We then benchmarked our medium A/SFT63 condition against a chemically defined (ChemD) culture medium, which has recently been advocated for the expansion of CB HSC (*23*). Importantly, ChemD was unable to expand mPB HSPC from n=2 adult donors (Fig.3G). Expansion of CD34+ cells was partially rescued when IL3 was added to ChemD confirming the importance of this cytokine for mPB cultures.

Third, we tested the addition of other small molecule compounds, which have been shown to promote HSC expansion, to the EX-U condition. CD34+ cells from mPB were transduced with the LVBC and expanded in medium A with SFT63 and UM171 (EX-U) or UM171 plus SR1, an aryl hydrocarbon receptor antagonist (EX-US) (Fig.3H). Xenotransplantation showed no statistically significant differences in human CD45+ cell engraftment of the EX-US condition compared to EX-U in the blood (Fig.3I) and the bone marrow (Fig.3K), and neither in terms of clonal graft diversity (Fig.3L; Table S4) or barcode sharing between mice (Fig.3M; Table S4) suggesting that EX-U and EX-US were similarly permissive to symmetric self-renewal divisions in culture. However, secondary transplantation showed significantly higher engraftment in the blood (Fig.3N) and a trend for higher bone marrow engraftment (Fig.3O) in the EX-US group. To better understand the dynamics of secondary repopulation, we analyzed the clonal abundance of shared clones between the primary and secondary grafts (Fig.3P; Table S5). In both EX-U and EX-US conditions, the major clones contributing to hematopoiesis in secondary mice were initially minor clones in the primary grafts, and vice versa, confirming that these assays are reading out different waves of hematopoiesis driven by LT-HSC with different degrees of latency. Thus, SR1, though not critical when using optimized culture conditions, may provide additional benefits in maintaining LT-HSC with latency characteristics in culture.

Fourth, we investigated the use of the DEGS1 inhibitor, 4HPR, in *ex vivo* expansion protocols. In line with previous reports (*21*), we confirmed that 4HPR increased the in vitro clonogenic output of CB HSC (CD34+CD38-CD90+CD45RA-CD49f+) but not of progenitors (CD34+49f-) (Fig.4A). Enhanced CFU forming potential was found also for sorted HSC from mPB when supplemented with 4HPR irrespective of UM171 addition (Fig.4B), also when 4HPR was added after LV transduction, during a 7-day *ex vivo* expansion period prior to CFU seeding (Fig.4C). Encouraged by these data suggesting a specific impact of 4HPR on mPB HSCs, we expanded another DP from an MPS patient (Pt5) with the triple combination of UM171, SR1 and 4HPR (EX-US4) and compared engraftment to EX-US, EX-U or the DP itself (Fig.4D). The EX-US4 condition showed significantly reduced BM engraftment as compared to EX-U at 12 weeks (Fig.4E), but no evident difference to the other conditions in secondary repopulation potential (Fig.4F). Clonal analysis by LVIS confirmed reduced number of unique insertion sites in the 4HPR supplemented culture (Fig.4G) with no major changes in clonal size distribution of primary grafts (Fig.4H). We confirmed no benefit also from the addition of 4HPR to UM171 alone in the absence of SR1 (EX-U vs EX-U4) for Pt1 and Pt2 xenografts (Fig.S3C,D).

**Figure 4:**
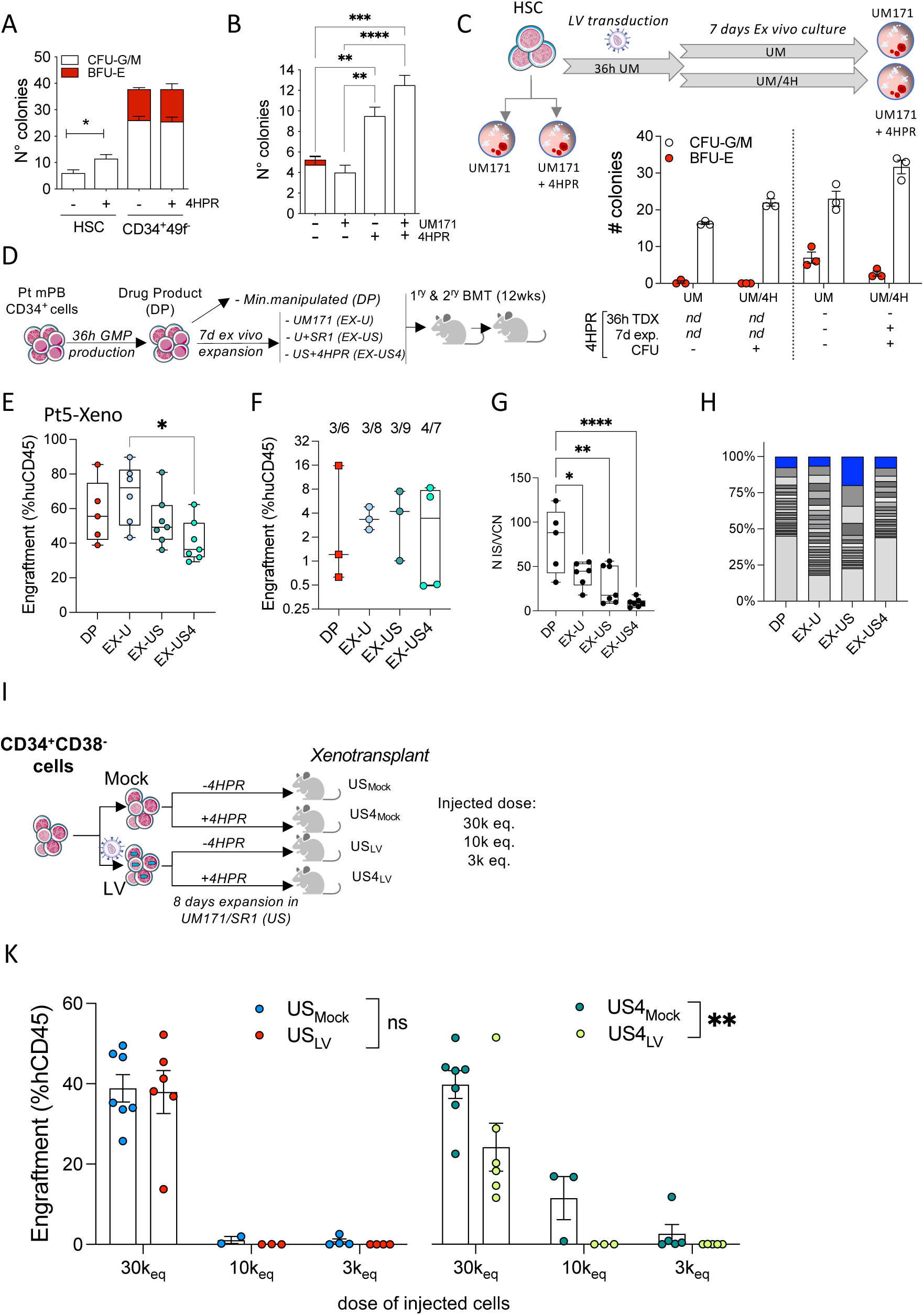
Cumulative toxicity between proteostatic cellular stress and lentiviral transduction. (**A**) Colony count in CFU assays with sorted HSC (CD34^+^CD38^-^CD90^+^CD45RA^-^CD49f^+^) or CD34^+^ (CD34^+^49f^-^) cells (pool of n = 4 CB donors) in presence (+) or absence (−) of 4HPR (2µM). The mean ± SEM of four replicates (600 cells/plate) is shown (unpaired t-test). (**B**) Colony count in CFU assays with sorted HSC (CD34^+^CD38^-^CD90^+^CD45RA^-^CD49f^+^) cells (n=1 mPB donor) in presence (+) or absence (−) of 4HPR (2µM) and or UM171 (35nM). The mean ± SEM of four replicates (800 cells/plate) is shown (One-way ANOVA with Bonferroni correction). (**C**) (Top) Experimental design. Expansion culture and/or colony assays were conducted with UM171 (UM) in presence (+) or absence (−) of 4HPR (4H). (Bottom) Colony count in CFU assays with sorted HSC (CD34^+^CD38^-^CD90^+^CD45RA^-^CD49f^+^) cells (n=1 new mPB donor) performed either on freshly sorted cells (2 groups on the left) or after lentiviral transduction and *ex vivo* expansion (2 groups on the right), in the presence (+) or absence (−) of 4HPR (2µM) as shown at the bottom of the graph. The mean ± SEM of three replicates (1000 cells/plate) is shown. (**D**) Schematic representation of xenotransplantation experiments performed on the DP from an MPS patient (Pt5). (**E**) Human cell engraftment in the BM of NSG mice at 12 weeks after transplantation of (i) 200,000 cells from the drug product (DP) (*n* = 5); (ii) 5,400,000 *ex vivo* expanded cells EX-U (equivalent to 200,000 CD34⁺ cells at day 0, expanded for 7 days with UM171; *n* = 6); (iii) 6,200,000 *ex vivo* expanded cells EX-US (equivalent to 200,000 CD34⁺ cells at day 0, expanded for 7 days with UM171 and SR1; *n* = 7); (iv) 1,300,000 *ex vivo* expanded cells EX-US4 (equivalent to 200,000 CD34⁺ cells at day 0, expanded for 7 days with UM171,SR1 and 4HPR; n = 7); Kruskal-Wallis test with Dunn’s multiple comparison, *p<0.05. (**F**) Human cell engraftment in the BM, 15 weeks after secondary transplantation of 1 × 10⁶ CD34^+^ cells per mouse, isolated from primary recipients described in (E). The number of mice with multilineage engraftment over the total number of transplanted mice is shown. (**G**) Box plots indicating the number of unique insertion sites normalized on vector copy number retrieved from the BM of primary grafts described in (E) One-way ANOVA with Tukey’s multiple comparison test. (**H**) Relative abundance of insertion sites in the primary recipient mice. Bottom light grey stacks indicate the sum of insertions with <1% relative abundance, middle stacks (shades of grey) indicate insertions >1% abundance, the top abundant insertions are highlighted in blue. (**I**) Schematic representation of xenograft experiment performed on the progeny of sorted CD34^+^CD38^-^ cells from adult mPB testing the following 2 variables: (i) transduction (LV) or not (Mock) with a purified lentiviral vector; (ii) expansion culture with or without 4HPR. (**K**) Human cell engraftment in the bone marrow was evaluated by analyzing the proportion of CD45⁺ cells 16 weeks post injection. Shown is the mean ± SEM, each dot represents an individual mouse, a mouse was classified as engrafted if it contained >0.1% of hCD45+ cells. ns, not significant; ** p<0.01 (Mock vs LV, Two-way Anova).

To understand whether LV transduction had a role in this unexpected outcome, we performed limiting dilution xenotransplantation experiments on mPB CD34+CD38- cells from day 8 expansion cultures with the EX-US or EX-US4 condition, following mock- or LV transduction (Fig.4I). Addition of 4HPR to UM171 and SR1 during *ex vivo* expansion gave similar engraftment in the absence of transduction but resulted in a significant loss of engraftment of LV transduced cells, especially at limiting cell doses (Fig.4K). A negative impact of LV transduction was also observed in the EX-US group, but only at limiting transplantation doses.

Taken together, these data suggest that LV transduction cumulatively adds to a stress response in the ex vivo expansion setting, which may render HSC functionally deficient in xenotransplantation assays. Combination of proteostasis inhibition with lentiviral transduction is thus not considered beneficial for mPB HSC gene therapy protocols.

### Controlling dose and transduction time window alleviates LV toxicity in the ex vivo expansion setting

Considering the unexpected sensitivity of *ex vivo* expansion cultures to LV transduction, we further investigated this critical genetic engineering step by shifting the LV exposure time window from the canonical time window of 24-48 hours after the start of *ex vivo* culture by 24 hours increments (Fig.5A). *In vitro* maintenance of CD34 expression was consistent across all conditions (Fig.5B), while longer pre-stimulation increased overall transduction efficiency (Fig.5C). However, engraftment potential of EX-U cells dropped sharply when transduction was carried out in the 72 to 96hr window (late transduction group, EX-U3), while it was equal or slightly superior to DP-like cells if transduction was conducted before (Fig.5D). Analysis of intragroup barcode sharing confirmed symmetric *ex vivo* HSC division, with a trend for less shared barcodes in the late transduction group compared to EX-U2 (Fig.5E, Table S6). To gain further insight into the mechanism behind LV-induced loss of HSC potency, we performed bulk RNA sequencing on *ex vivo* expanded HSPCs transduced at early (24hr) or late (72hr) timepoints. CD34+ HSPC exposed to the LV during the 72-96hr window of *ex vivo* expansion showed strong enrichment of senescence- and aging-associated pathways (Table S7, Fig.5F). Based on these findings, we selected an intermediate time window from 36 to 48hr (EX-U2) after starting pre-stimulation, as the optimal time for LV transduction in an *ex vivo* expansion setting, which preserves HSC engraftment potential.

**Figure 5:**
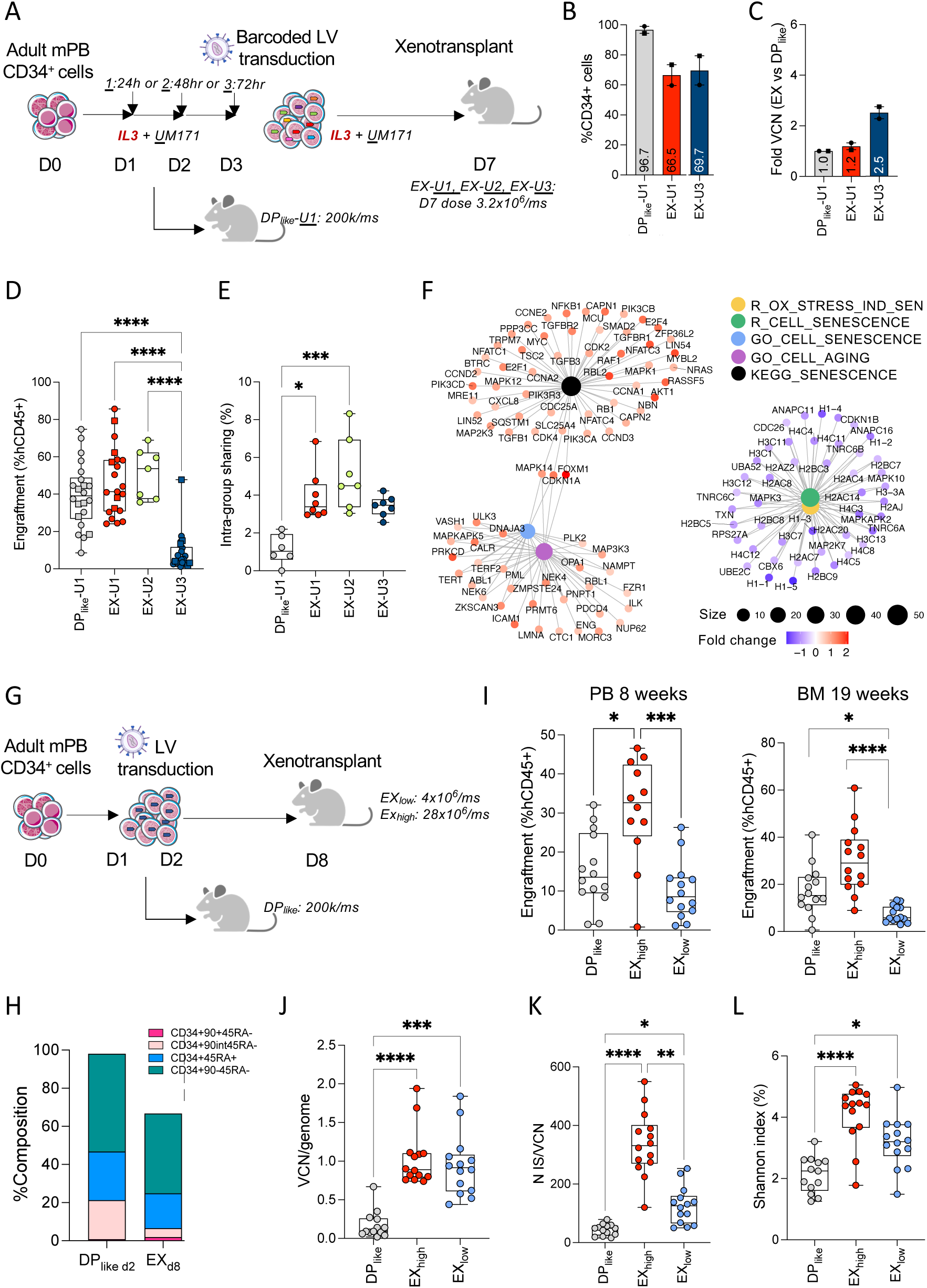
Controlling dose and transduction time window alleviates LV toxicity in *ex vivo* expansion setting. (**A**) Schematic representation of the experimental design for data shown in panels B-F. Two separate transplant sessions were performed using CD34+ cells from the same adult mPB donor under the same protocol, each session transplanted cell products from all groups, except the EX-U2 condition, which was added during the second transplant session. Pooled results from the 2 sessions are shown. (**B**) The percentage of CD34^+^ cells measured by flow cytometry after 7 days of expansion under different conditions. Bars represent the average from the 2 experimental sessions. Data not shown for EX-U2 (76%) as only a single replicate was performed. (**C**) Relative change in the vector copy number per genome (VCN) in the EX-U condition with respect to DP_like_, as determined by ddPCR on liquid culture (n=2 experimental sessions; n=1 for EX-U2 [4.8] data not shown). (**D**) Engraftment potential of HD mPB CD34^+^ hematopoietic cells under different conditions, by transplantation of NSG mice with either: (i) 200,000 cells from the DP_like_-U1 condition (*n* = 22), (ii) 3,200,000 *ex vivo* expanded cells (equivalent to 200,000 CD34⁺ cells at day 0, expanded for 7 days and transduced at the following time points: EX-U1, 24h, n=21; EX-U2, 48h, n=7; EX-U3, 72h, n=22. Human cell engraftment in the bone marrow was evaluated by analyzing the proportion of CD45⁺ cells 17 weeks post injection (Kruskal-Wallis test with Dunn’s multiple comparison). (**E**) Intra-group sharing percentages assessed with shared LVBCs (HD1). Each dot represents the percentage of sharing between an individual mouse vs all other mice receiving the same HSPC product (Kruskal-Wallis test with Dunn’s multiple comparison, *p<0.05, ***p<0.001, **** p< 0.0001). (**F**) Bulk RNA sequencing was performed to identify DEGs between the EX-U3 and EX-U1 groups. The CNET plot shows enriched senescence terms obtained from GSEA, blue and red dot colors represent down/up regulated genes, respectively. The dots size represents the number of genes that belong to the term. (**G**) Experimental scheme. Adult mPB CD34+ cells were transduced with a clinical grade LV and either expanded (EX) or not (DP_like_) using a GMP-compliant manufacturing process. The following groups were injected into NSG mice: (i) 200,000 DP_like_ cells (*n*=14), (ii) 20,000,000 *ex vivo* expanded cells at high dose EX_high_ (equivalent to 530,000 CD34⁺ cells at day 0; *n*=12) or (iii) 4,000,000 *ex vivo* expanded cells low dose EX_low_ (equivalent to 100,000 CD34⁺ cells at day 0; *n*=14). (**H**) Percentage of CD34^+^ cells and subset composition measured by flow cytometry in the respective products (day 2, DP_like_; day 8, EX) before xenotransplantation. (**I**) Human CD45+ cell engraftment measured in the peripheral blood (left) and bone marrow (right) at the indicated timepoints (Kruskal-Wallis test with Dunn’s multiple comparison). (**J**) Vector copy number (VCN) per genome determined by ddPCR on BM cells from mice of different groups (Kruskal-Wallis test with Dunn’s multiple comparison). (**K**) Box plots indicating number of unique insertion sites normalized on vector copy number, (Kruskal-Wallis test with Dunn’s multiple comparison). Each dot represents an individual mouse of the treatment group: DP_like_, EX_High_ and EX_low_ (Kruskal-Wallis test with Dunn’s multiple comparison). (**L**) Box plots indicating the Shannon diversity index as measure of clonal population diversity, (Kruskal-Wallis test with Dunn’s multiple comparison). Each dot indicates an individual mouse of the treatment group: DP_like_, EX_High_ and EX_low_.

Finally, a putative drug product manufactured according to our optimized expansion was compared to a transduction protocol (EX) with a clinically validated short manufacturing protocol (DP_like_) for gene therapy (Fig.5G,H) (*30, 32*). The comparison was conducted using a GMP-compliant, large-scale process run in gas-permeable bags and a therapeutically relevant, purified lentiviral vector, developed for the treatment of autosomal recessive osteopetrosis, a setup suitable for clinical translation (*32*). Engraftment of DP_like_ (2×10^5^ cells/mouse) was compared to a low or high dose of EX (EX_low_: 4×10^6^ and EX_high_: 28×10^6^ cells/mouse, corresponding to a d0 equivalent of 1×10^5^ and 7×10^5^ cells/mouse, respectively; n=14 per cohort). Multi-lineage human cell engraftment in the blood at 8 weeks post transplantation and in the BM at 19 weeks (Fig.5I) was highest for the high-dose EX_high_ group, while the DP_like_ group positioned in between high- and low-dose EX_low_. These data align with a net HSC maintenance during *ex vivo* culture when considering the overall graft. However, engraftment of genetically engineered cells, which are the therapeutically relevant part of the graft, was higher in EX as compared to DP_like_, even in the low-dose EX_low_ group, as shown by an increased number of vector copies (VCN) per human genome (Fig.5J), an increased absolute number of unique integration sites normalized to VCN (Fig.5K) and an increased Shannon diversity index (Fig. 5L). These results highlight an advantage of the optimized expansion protocol over the current standard, even at lower transplant cell doses.

## DISCUSSION

Efforts to optimize *ex vivo* HSC manipulation protocols have mainly focused on minimizing culture time to preserve repopulating potential, reduce clonal loss, and mitigate senescence or genomic instability. The inclusion of molecules that antagonize undesired culture effects and the careful protocol optimization we have undertaken have made it possible to extend *ex vivo* culture time of mPB to at least a week, without losing HSC function or clonality compared to state-of-the art short culture protocols. Not surprisingly, the degree of net HSC expansion was lower in mPB as compared to published CB studies (*12, 18*), which may be explained by higher functional HSC content (*33*) and faster cell cycle kinetics (*34*) in the latter. Nevertheless, expanded mPB products exhibited potent multilineage engraftment potential, higher vector copy numbers per genome, and preserved clonal diversity even at lower transplant doses compared to shortly cultured cells. Thus, *ex vivo* expansion of mPB HSCs represents a promising avenue for overcoming limitations in cell availability and enhancing therapeutic efficacy for gene therapy applications.

We present the stepwise optimization of a clinically translatable, GMP-compliant expansion protocol tailored for mPB-derived HSCs from both adult and pediatric donors. We optimized cytokine combinations by exploiting predicted receptor expression in scRNAseq identifying a benefit in adding intermediate doses of interleukin-3 to promote the growth of progenitors while maintaining the absolute number of phenotypic HSC, potentially by stimulating asymmetric HSC division. Even though the receptor for interleukin-6 does not appear to be highly expressed on HSC, intermediate doses of IL-6 increased the number of phenotypic HSC. This may potentially be due to non-cell autonomous effects, e.g. by stimulating crosstalk between monocyte precursors, on which the IL-6 receptor is upregulated, and HSCs, a phenomenon that has been described to enhance engraftment potential (*35*). Often overlooked in protocol design, media selection significantly influenced the yield of functional HSCs as defined by xenograft assays, a cumbersome readout generally not performed by developers of commercial cell culture media. Lack of transparency on formulations and single ingredients preclude identification of the beneficial and harmful components contained in the different commercial products. Lastly, we explored combinations of HSC maintaining molecules on top of UM171, which in our hands (*29*) and in the hands of others (*12, 23*) has been the most potent compound. Our data now show an added benefit of blocking the aryl hydrocarbon receptor pathway by SR1, most notably on ST-HSC, but also on LT-HSC when challenged in secondary repopulation assays or under non-optimal *ex vivo* expansion conditions (data not shown). We observed toxicity from the addition of 4HPR to mPB HSCs in our protocol, aggravated by LV transduction. While the exact mechanism of this toxicity remains to be investigated, we may speculate that mPB HSCs, by themselves poised to an increased basal pro-inflammatory state (*36, 37*), which is further enhanced by cytokine pre-stimulation and LV exposure, may exhaust compensatory mechanisms when challenged with 4HPR-induced oxidative/ER stress (*21*).

An unexpected finding was also the critical influence of the timing of LV transduction on HSC functionality. The response of human HSPC to exposure of purified LVs has been described as rather limited, with some degree of p53 activation through nuclear sensing of vector DNA but little evidence for innate immune activation (*31*). This is in line with clinical data whereby even highly transduced DPs engrafted promptly and did not cause measurable side effects, in the context of a short *ex vivo* culture protocol (*30*). Interestingly, the sensitive time window of LV transduction coincides with the first cell division of purified mPB LT-HSC, estimated to occur around 62.0 ± 6.5 hours (*11*), which may increase the susceptibility of HSC to undergo senescence upon p53 activation. These findings require special attention when performing prolonged *ex vivo* HSC culture and may have broader implications for HSC manipulations beyond LV transduction. Notably, a recent study has found a very similar vulnerability time window of HSC to gene editing triggering a DNA damage response, which could be mitigated by slowing cell cycle progression through p38 MAPK inhibition (*38*).

A major challenge in successful development of *ex vivo* expansion protocols is accurately predicting in vivo repopulating capacity based on in vitro product characteristics. Many validated surface HSC marker profiles, such as CD38, lose their predictive value once cells are manipulated *ex vivo*, or they may not be consistently expressed across cell sources (*29, 39, 40*). We thus undertook an unbiased characterization of *ex vivo* cultured HSPC by scRNAseq, which provided high-resolution insights into cellular heterogeneity and lineage fidelity of committed precursors during culture. Additionally, scRNA-seq enabled predictive analyses confined to functionally relevant, select cell populations, namely molecularly defined HSC, in which *in vitro* VCN prediction correlated with the VCN measured in xenografts and, importantly, with the VCN reported in patients following long-term engraftment (*30*). Expansion culture was required to accurately assess transduction efficiency in HSCs, since direct analysis of the DP may yield false positive results, presumably due to the presence of transcripts from non-integrated vector forms. Hence, this provides a proof of concept that ex vivo expansion cultures may serve as a cost-effective surrogate for *in vivo* cell product dynamics assessment, that may eventually replace xenograft experiments, even though additional validation on a higher number of samples is needed.

Clonal tracking analyses stringently validated our *ex vivo* culture by assessing for the presence of symmetric self-renewal divisions *ex vivo* with increased sharing of genetic marks across mice transplanted from expansion culture as well as ruling out clonal dominance or oligo-clonal selection—a critical safety consideration in gene therapy applications. A recent study has used barcoding to show self-renewal of molecularly defined HSC (by scRNAseq) during *in vitro* culture of CB in the presence of UM171 (*41*). We add to this conclusion by demonstrating self-renewal of functionally defined HSC derived from mPB. Integrating scRNAseq-based identification of HSC transcriptional states with *in vivo* clonal tracking by scRNAseq-compatible barcoded LV libraries, could yield even deeper insights into cellular states and dynamics during *ex vivo* expansion and related *in vivo* implications. Our clonal bulk analysis *in vivo* revealed distinct waves of hematopoietic repopulation between primary and secondary xenografts, mirroring what has been previously described as the output of latent LT-HSC subsets resistant to minor stress-induced activation (*42*). These latent CD112^lo^ LT-HSCs are supposed to preserve self-renewal capacity by remaining quiescent under stress conditions but are recruited during major regenerative demands such as those imposed by a secondary transplantation. Nevertheless, our data suggest that these latent HSC have a memory of the *ex vivo* culture, as suggested by improved secondary reconstitution of HSC exposed to the combination of SR1 and UM171. The ability to show polyclonal engraftment in secondary transplants with our protocol is again a strong indicator of preservation of bona fide HSC functionality with our optimized expansion protocol.

Despite our in-depth functional and molecular characterization of *ex vivo* mPB expansion cultures down to single cell/single clone resolution, and systematic process optimization, we could not obtain a consistently high expansion of mPB LT-HSC numbers, which may in part be explained by distinct biological properties with respect to CB. Two approaches can be envisioned to further increase net HSC expansion: (1) Stimulate HSC self-renewal divisions, by targeted molecular interventions in the self-renewal program (*43*) or, most simply, by increasing the length of culture considering that slow cell cycle entry and progression is a core property of LT-HSC (*44*). (2) Mitigate the loss of HSC through irreversible differentiation, e.g. by reducing culture stress through inhibition of pathways such as JAK (*11*), MAPK (*38*) or mTORC (*10*), titration of oxygen levels (*45*) and mechanoregulation (*46*). It should be highlighted that our functional readouts concentrate on HSC and do not assess early hematopoietic reconstitution driven by progenitors, which cannot be read out in the NSG model and has not been further evaluated in this work. Hence, *ex vivo* mPB expansion with our optimized protocol is expected to offer clinical benefit in terms of myeloid and platelet reconstitution over non-cultured or minimally cultured cells, pending confirmation in future clinical trials. Notably, we have submitted a clinical trial application for a phase I/IIa gene therapy study (EU CT 2024-518972-30) in children affected by autosomal recessive osteopetrosis (ARO), a rare and severe genetic bone disorder characterized by defective osteoclast function and the lack of a BM HSC niche, where rapid reconstitution of resorption-competent osteoclasts may allow proficient engraftment of corrected LT-HSC (*47*). In sum, we have developed an optimized *ex vivo* mPB expansion protocol to aid in genetic correction of *ex vivo* expanded autologous circulating HSCs to address unmet medical needs (*32*).

## MATERIALS AND METHODS

### Human Hematopoietic Stem and Progenitor Cells

Human CB and mPB CD34+ HSCs were purchased from commercial sources (Lonza, HemaCare, All Cells and Charles River). Aliquots of drug products from patients enrolled in the TigetT10_MPSIH trial (NCT03488394, see Gentner et al, N Engl J Med 2021 (*30*) for relevant design details) and specifically allocated for research purposes were available in accordance with the clinical trial protocol and the associated research protocols (TIGET09), which have been approved by the responsible ethical committees. Informed consent for the use of these materials in research was obtained from all patients’ legal guardians.

### Sample size and replicates

Sample size pre-determination was used for the experiment validating the optimized expansion culture setup (Fig. 5G-L). For the experiments with patient cells, exploratory experiments and iterative process optimization steps, sample size for each experiment was dependent on the total number of available HSPCs, which is constrained by the human source of the material and limited by transduction efficiency, scale, and recovery from cell expansion culture or primary recipient xenografts. Whenever possible, we aimed to reach n ≥ 5 per group, which is considered adequate for carrying out nonparametric statistical comparisons.

Data was collected at prespecified endpoints for transplantation experiments, with primary recipients euthanized at 12 weeks post engraftment if secondary transplants were to be carried out, or at longer timepoints (≥16 weeks). All samples passing prespecified quality control metrics were included in the presented analyses. No outliers were excluded.

Due to the limitations on animal experimentation, some exploratory experiments were performed once. If replicate experiments have been conducted, they are specified within the specific figure legend.

### Study design and research objective

We here show data from controlled laboratory experiments. Each experimental design is appropriately outlined in the text or figure and addresses a specific hypothesis for the improvement of *ex vivo* culture protocols for mobilized peripheral blood HSPC expansion. When performing mouse experiments, mice were randomly allocated to the control or experimental arm. Experiments were not conducted in a blinded fashion, as blinding was not relevant for objective outcome measures.

### HSPC culture, transduction and expansion protocols

HSPC pre-stimulation and transduction was performed in serum-free medium, at a cell density of 1×10^6 cells/ml. For the *ex vivo* expansion culture, cell density was reduced to 2×10^5 cells/ml and maintained at <1×10^6 cells/ml. Specific culture media and cytokine addition varied across experimental plan, as detailed in the results section and figure legends.

Mobilized peripheral blood HSPC drug products (DP) were cultured in SCGM medium (Cell Genix) supplemented with the following cytokines: 60 ng/ml IL-3, 100 ng/ml TPO, 300 ng/ml SCF, and 300 ng/ml FLT-3L (all from CellGenix) (*48, 49*).

Expansion cultures were set up in serum-free medium (SCGM CellGenix except where differently indicated) containing, unless otherwise indicated, 100 ng/mL SCF, 100 ng/mL FLT3L, 50 ng/mL TPO, 50 ng/mL IL-6, 35 nM UM171 (Stem Cell Technologies or ExCellThera) and (±SR1 750nM; StemRegenin1 from Stem Cell Technologies) (±4HPR 2µM; kind gift from the J.E. Dick laboratory). Cultures were kept at 37°C, 5% CO_2_ under normoxia. A chemically defined expansion medium was adapted from (*23*) and https://doi.org/10.1101/2024.09.17.613552, as follows: Iscove’s Modified Dulbecco’s Medium (IMDM) supplemented with Soluplus (BASF) 1mg/ml, 740 Y-P (Cayman Chemical) 1µM, Butyzamide (TargetMol) 0.1µM, monothioglycerol (Sigma) 100µM, UM729 (Stem Cell Technologies) 1µM, ITSX, Glutamax and low dose SCF and FLT3L (CellGenix), both 25ng/mL. LV transduction was performed by addition of 10^8 transducing units/ml (multiplicity of infection: 100, unless otherwise indicated) to the cell cultures for 16-24 hours.

### LV Production, Titration, and Molecular Analysis of Gene Transfer Efficiency

Purified vectors (PGK.IDUA and TCIRG1 LV) were produced and titered by MolMed (now AGC Biologics), according to the GMP process employed for gene therapy clinical trials (*48, 49*). Lab-grade vectors, third generation self-inactivating (SIN) LV (PGK.GFP, PGK.BFP, PGK.mCherry) expressing different fluorescent protein under the control of a PGK promoter stocks were produced and titered on 293T cells according to standard lab protocols as previously described (*50*). Large-scale LV barcode library was produced and purified by a specialized institutional process development lab as described in Soldi et al. (*51*) Molecular quantification of gene transfer efficiency was performed using specific ddPCR assays.

### Clonogenic assay on Methocult

Clonogenic assays were plated at the time point described for each single experiment. Cells were washed, counted and resuspended in complete human Methocult medium (Stem Cell Technologies) at a concentration of 800-2000 cells/ml. Fourteen days later, colonies were scored by light microscopy for number and morphology. CFU-E and BFU-E were scored as erythroid colonies, while CFU-G, CFU-M and CFU-GM as myeloid and CFU-GEMM as mixed colonies. If necessary, at day 14, colonies were single-picked and analyzed by ddPCR to evaluate transduction efficiency.

### Determination of VCN by droplet digital PCR

Genomic (g)DNA was extracted using QIAmp DNA micro kit (Qiagen) or QIAmp DNA mini kit according to the starting number of cells (as suggested by manufacturer). DNA was quantified and assessed for purity. Vector copies per diploid genome (vector copy number, VCN) of the integrated lentiviral vectors were quantified by digital droplet PCR (ddPCR) on a QX200 Droplet Digital PCR System (Bio-Rad) starting from 10-100 ng of template gDNA using the following primers (HIV sense: 5’-TACTGACGCTCTCGCACC-3’; HIV antisense: 5’-TCTCGACGCAGGACTCG-3’) and probe (FAM 5’-ATCTCTCTCCTTCTAGCCTC-3’) against the primer binding site region of LVs. Endogenous DNA amount was quantified by a primer/probe set against the human telomerase gene (Telo sense: 5’-GGCACACGTGGCTTTTCG-3’; Telo antisense: 5’-GGTGAACCTCGTAAGTTTATGCAA-3’; Telo probe: VIC 5’-TCAGGACGTCGAGTGGACACGGTG-3’ TAMRA). Alternatively, region of the primate transcription initiation factor TFIID subunit 7 gene (TAF7), which is conserved among primates and humans, was used as an endogenous control assay (sequence not disclosed). The VCN was determined by calculating the ratio of the target molecule concentration to the reference molecule concentration, times the number of copies of reference species in the genome. All the reactions were performed according to the manufacturer’s instructions and analyzed with a QX200 Droplet Digital PCR System (software: QuantaSoft Version1.7.4.0917; Bio-Rad).

### Cell Sorting and Flow Cytometry

Sorting for CD34+ cells (and CD38 cells, where indicated) was performed by magnetic beads (Miltenyi). Fluorescence-activated cell sorting for HSPC subpopulations was performed on a MoFlo XDP sorter (Beckman Coulter) or BD FACSAria Fusion (BD). Analytical flow cytometry was performed on a FACSCanto II or LSR II Fortessa instrument (BD Bioscience).

Immunophenotypic analyses were performed by flow cytometry using Canto II (BD Pharmingen). From 5.0 × 10^5 to 2.0 × 10^6 cells either from culture or mouse-derived samples were analyzed. Cells were stained for 15 min at 4 °C with antibodies listed in Table 1, in a final volume of 100μl and were washed with DPBS + 2% heat inactivated FBS. Single-stained and fluorescence-minus-one-stained cells were used as controls.

**Table 1.** List of anti-human antibodies used for flow cytometry.

| Antibody | Fluorochrome | Clone | Company | Code |
| --- | --- | --- | --- | --- |
| FcR Blocking Reagent | - | - | Miltenyi Biotec | 130-059-901 |
| CD45 | APC-eFluor780 | HI30 | eBioscience | 47-0459-42 |
| CD45 | VioBlue | clone REA747 | Miltenyi | 170-081-077 |
| CD19 | PE-Cy7 | HIB19 | BioLegend | 302216 |
| CD19 | PE | clone SJ25C1 | BD Biosciences | 345789 |
| CD3 | PE | SK7 | BD Biosciences | 345765 |
| CD3 | PB | UCHT1 | BioLegend | 300431 |
| CD13 | APC | WM15 | BD Biosciences | 557454 |
| CD33 | APC | AC104.3E3 | Miltenyi Biotec | 130-091-731 |
| CD33 | PE-Cy7 | P67.6 | BD Biosciences | 333952 |
| CD33 | APC-Vio770 | clone REA775 | Miltenyi Biotec | 130-111-022 |
| CD34 | VioBlue | AC136 | Miltenyi Biotec | 130-095-393 |
| CD34 | PE-Cy7 | clone 8G12 | BD Biosciences | 348811 |
| CD34 | APC | AC136 | Miltenyi Biotec | 130-120-519 |
| CD38 | PE-Vio770 | IB6 | Miltenyi Biotec | 130-099-151 |
| CD90 | APC | 5E10 | BD Biosciences | 559869 |
| CD90 | BV421 | 5E10 | BD Biosciences | 562556 |
| CD45RA | PE | T6D11 | Miltenyi Biotec | 130-092-248 |
| CD45RA | FITC | clone HI100 | BioLegend | 304106 |
| CD56 | PE | clone NCAM 16.2 | BD Biosciences | 345812 |
| BAH1 | PE | BAH-1 | BD Biosciences | 565746 |
| CD71 | BV711 | M-A712 | BD Biosciences | 563767 |
| CD49f | PC5 | GoH3 | BD Biosciences | 562495 |
| 7AAD | - | - | Miltenyi Biotec | 130-111-568 |

### Mice

All experiments and procedures involving animals were performed with the approval of the Animal Care and Use Committee of the San Raffaele Hospital (IACUC 923, 1183, 1207) and were authorized by the Italian Ministry of Health and local authorities accordingly to Italian law. NOD-scid-Il2rg−/− (NSG) female mice (The Jackson Laboratory) were held under specific pathogen-free conditions.

### CD34+ HSPC xenotransplantation experiments in NSG mice

For xenotransplantation of mPB HSPCs, the outgrowth of 2×10^5 to 5×10^5 HSPCs at the start of the culture (d0 equivalent) were injected intravenously into sublethally irradiated (180–200 cGy) female NSG mice. Matched numbers of HSPCs were seeded at day 0 of culture for each experimental group to transplant the same number of culture-initiating HSPCs in each mouse. The precise dose of injected cells differed according to the experimental conditions and is described in the figure legend for each experiment. Mice were randomly distributed to each experimental group. Human CD45+ cell engraftment and the presence of transduced cells were monitored by serial collection of peripheral blood (approximately every 4 weeks) and, at the end of the experiment (12–18 weeks after transplantation), BM was collected for end point analyses. Hind limb BM was flushed with PBS 2% FBS and 2×10^6 cells were stained for surface markers. The remaining cells if required were used to performed secondary transplantation in NSG mice after beads purification with mouse cell-depletion or enriched for human CD34+ cells kit (Miltenyi Biotec) according to the manufacturer’s instructions.

### Peripheral blood analysis

Mice were bled via the tail vein following analgesia. For each mouse, 250μl of peripheral blood was added to 10μL of PBS containing 45mg/mL EDTA. For immunostaining, a known volume of whole blood (100μl) was incubated with antihuman Fc receptor blocking antibodies for 10 min at room temperature and then incubated in the presence of monoclonal antibodies (for a list of antibodies, see Table 1 above) for 15 min at room temperature. Erythrocytes were removed by lysis with the TQ-Prep workstation (Beckman-Coulter) in the presence of an equal volume of FBS (100μl).

### Bone marrow analysis

At the experimental endpoint, mice were humanely euthanized, and BM cells were obtained by flushing the femurs in PBS 2% FBS solution. Cells (1-2×10^6 cells) were washed, resuspended in 100μl of PBS containing 2% FBS, and incubated with anti-human and/or anti-mouse FcγIII/II receptor (Cd16/Cd32) blocking antibodies for 15 min at 4°C. Staining was performed with monoclonal antibodies (Table 1 above) for 20 min at 4°C.

### Lentiviral Barcode NGS Library Preparation

PCR products compatible with Illumina sequencing were generated by targeted amplification of the integrated lentiviral vector barcode from genomic DNA (gDNA) isolated from the bone marrow of murine xenografts. A total of 50–100 ng of gDNA was used as input. Amplification was performed using Phusion High-Fidelity DNA Polymerase (Thermo Fisher Scientific) according to the manufacturer’s protocol, with the following thermal cycling conditions: initial denaturation at 98°C for 60 seconds; 26 cycles of denaturation at 98°C for 10 seconds, annealing at 56°C for 30 seconds, and extension at 72°C for 15 seconds; followed by a final extension at 72°C for 10 minutes.

Custom next-generation sequencing (NGS)-grade primers were used at a final concentration of 10 μM. Primers included Illumina adapter sequences (P5 and Read 1 for forward primers; P7 and Read 2 for reverse primers) along with unique 8-base sample-specific barcodes (denoted as “xxxxxxxx”) for multiplexing.

Primer sequences were as follows:

- Forward primer: 5’-AATGATACGGCGACCACCGAGATCTACACTCTTTCCCTACACGACGCTCTT CCGATCTxxxxxxxxCTACTCAGACAATGCGATGC-3’
- Reverse primer: 5’-CAAGCAGAAGACGGCATACGAGATGTGACTGGAGTTCAGACGTGTGCTCT TCCGATCTxxxxxxxxCGTGCCTTCCTTGACCCTG-3’

PCR products were resolved on a 1.35% agarose gel, and the 255 bp band corresponding to the expected barcode amplicon was excised and purified using the Promega Wizard® SV Gel and PCR Clean-Up System in accordance with the manufacturer’s instructions. Purified libraries were pooled at equimolar concentrations and subjected to paired-end sequencing (2 × 75 bp) on an Illumina NextSeq platform.

### Lentiviral Integration site library preparation

Integration sites were retrieved by Sonication Linker Mediated (SLiM)-PCR, as previously described (*52*), with minor modifications. Briefly, for each sample up to 300 ng of gDNA were processed by DNA shearing using the Covaris E220 Ultrasonicator, generating fragments with an average size of 1000 bp. The fragmented DNA samples were subjected to end repair and 3’ adenylation and then ligated to linker cassettes containing an 8-nucleotide sequence barcode used for sample identification, and all the sequences required for the Read 2 paired end sequencing. The ligation products were split in three technical replicates and subjected to 35 cycles of exponential PCR using primers specific for the lentiviral vector LTR and the Linker cassette. A subsequent amplification with additional 10 PCR cycles was performed using a primer specific for the linker cassette and the LTR. These primers contain an 8-nucleotide length barcode used for sample identification (coupled with the barcode on the linker cassette) and the sequences needed for the Read 1 sequencing, plus a 12 random nucleotides sequence allowing easier cluster recognition in the first sequencing cycles on the Next Generation Sequencing (NGS) sequencer. The generated SLiM-PCR products are thus associated to a unique pair of barcodes, assembled into libraries and subjected to NGS Illumina sequencing.

### LVIS bioinformatic analysis

Sequencing reads were processed using VISPA2 (*53*) that isolates genomic sequences flanking the vector LTR and maps them to the -human genome (hg19). Because vector integration in the same genomic position in different cells is a very low probability event, identical IS in independent samples were considered as contamination or amplification artefacts, which may occur during the technical procedure. Datasets were pruned from potential contaminations and false positives between each primary mouse and from IS deriving from unrelated secondary mice. - IS with identical genomic coordinates shared between mice belonging to different experimental groups, were reassigned based on identification of the insertion site by at least two SLiM technical replicates and sequence count numbers. Downstream analyses of vector integration sites, such as relative abundance analysis, sharing analysis and CIS analysis, were performed using ISAnalytics (*54*). Clonal abundance estimates as the relative percentage of genome numbers was determined by the R package “sonicLength” (*55*). Common insertion sites were calculated by the Grubbs test for outliers (*56*)

### Lentiviral barcoding preprocessing and quantification

Demultiplexed raw data were quality checked using fastqc tool to check for low-quality samples in terms of base quality scores and to check the presence of a barcoded region characterized by high heterogeneity. All samples were down sampled to 300k input reads using seqtk (1.4-r130) to avoid coverage biases and allow direct comparison of all runs and conditions, as well ensure reproducibility using a fixed seed. The distribution of barcodes counts, and Shannon entropy indexes were compared between raw and subsampled samples to verify that complexity and library saturation were maintained Identification and quantification of barcodes was performed with barseq (*57*) setting edit distance and minimal counts to 2 and 3 respectively. Barcodes extraction parametrization was the following: −1 P:CTACTCAGACAATGCGATGC −2 S:AATTTCCTCATTTTATT −3 R:N −4 S:TACGTCGA −5 P:GCGAGGA. Extraction metrics provided by TagDust v2.33 were hence inspected to identify and discard samples with low extraction rates (< 80%).

### Lentiviral barcoding processing and downstream analysis

To assess the threshold for the minimum barcode abundance to identify a clone we employed a strategy that relies on the comparison of shared clones between 1^ary^ and 2^ary^ transplant samples. We leveraged SCGM(CG) barcoded libraries treated with UM171 (UM) and UM171+SR1 (US) for which primary and secondary transplant data were available (Data S4). Using raw barcode counts provided by barseq, we looked for the lowest frequency barcode in 1^st^ transplant that was expanded in 2^nd^ transplant. Barcodes present in more than 3 mice in 1^st^ transplant group were considered artifacts and discarded. The identified threshold equal to (0.005%) was then applied to all the experimental groups analyzed in this work. Barcodes with frequency less than 0.005% were considered artifacts, contaminants or unreliable.

Moreover, we applied two strategies to further discard or minimize contaminants. The first strategy aimed to discard barcodes that were shared in more than N animals (Nmax parameter, see code for details) within each experimental group, whereas the second one consisted in the re-assignment of barcodes that were shared between independent groups. In particular, the barcode was re-assigned to the top sample whose quantification is 10X higher than the 2nd most abundant sample sharing the barcode. If none of the samples meet this rule, the barcode was considered shared across samples of independent groups.

We focused on the evaluation of the barcodes sharing within and between groups of samples as well as the clonal distribution and complexity. This latter topic was assessed evaluating the Shannon entropy index using the vegan R package (v.2.5-7).

Barcode count data processing and figure preparation was performed within R environment (v4.1.0), using several packages including among the most used: ggplot2 (v3.3.5), ggalluvial (v0.12.3) and dplyr (v1.0.8).

### Bulk RNA sequencing

Raw data in fastq format were QC checked and trimmed using TrimGalore (v0.5.0) to get rid of adapters at 3’ of reads. Trimmed reads were then aligned to the GRCh38 reference genome using STAR (v2.7.0d). Gencode genes primary assembly gene transfer file (GTF) v35 was used as reference gene annotation file. Post-alignment metrics, including coverage distribution across gene length and percentage of reads mapping to exons were collected by using Qorts (v1.3.5). We then assigned reads to genes (gene counting) by using feature counts (v1.6.3). Gene counts matrices were analyzed in the R environment (v4.0.3). We employed a data analysis workflow that relied on edgeR DEGs identification setting custom biological variation coefficients (BCV) to 0.1-0.3 for the different 1vs1 comparisons performed (72h vs 24h – in each batch separately). Counts were normalized using TMM method and differential test was performed with exactTest function provided by the edgeR package (v3.32). Genes with an adjusted p.value (Benjiamini Hochberg FDR method) < 0.05 were considered differentially expressed.

DEGs lists and pre-ranked gene lists according to log2FC were used for performing Over-Representation Analysis (ORA) and Gene Set Enrichment Analysis (GSEA) respectively. Terms with a p.adjust < 0.05 were considered statistically significant. Analyses and charts were produced with clusterProfiler (v4.7.1) (*58*), enrichplot (v1.16.2) and ggplot2 (v3.3.5) R packages.

GSEA pre-ranked list for the evaluation of the transduction timing was prepared calculating a mean fold change value of the two batchesadding an offset of 1 to avoid infinite values in log transformation. Both ORA and GSEA were computed considering different reference datasets including Gene Ontology, KEGG pathway database, Reactome pathway database and Molecular Signatures Database (MSigDB). In addition, for the evaluation of senescence signatures, we relied on a custom gmt reference file that includes literature and database-based signatures.

### Single-cell RNA sequencing

HSPCs were resuspended at the appropriate concentration for loading into the Chromium 10X single-cell 3′ Gene Expression v2 or v3 chip according to the manufacturer’s protocol. 3′ gene expression libraries construction and sequencing on Illumina platforms NextSeq or NovaSeq S1 were performed following the manufacturer’s indications.

Raw data were processed by Cell Ranger Single-Cell Software Suite, with cellranger count pipeline (version 7.0.1, 10X Genomics) to produce feature-barcodes matrices with Unique Molecular Identifier (UMI) counts for each cell. GRCh38 genome genes annotations provided by the manufacturer were used as reference data. Feature barcode counts matrices were then imported and processed within the R environment (v4.1.3) and analyzed with several R and Bioconductor packages. Fig.2C and Fig.2H datasets were analyzed with Seurat R package (*59*). Fig.2H dataset raw matrix was pre-processed with Adaptively-thresholded Low Rank Approximation (ALRA) algorithm using default settings. For both datasets we performed quality control by discarding cells with lower number of expressed genes and genes expressed by few cells. Counts were log normalized with a scale factor of 10,000 and most variable genes were identified (the top 15% of expressed genes). Normalized data was then scaled accounting for cell cycle (CC.Difference), total depth (nCount_RNA) and mitochondrial content (pct_mito). Hence, we performed dimensionality reduction by calculating principal components (PCs) and selecting the top components based on the elbow rule. Harmony methods were employed to remove potential batch effects, using sample identity or Timepoint + Patient (pt.tp) as the batch variable for Fig.2C and Fig.2H datasets respectively. Clustering was computed by Louvain improved algorithm on neighborhood graph built on top harmony components. Marker genes for each cluster were computed using the FindAllMarker function in Seurat package. Parameters used for each dataset can be found in the GitLab repository provided with the manuscript. For Fig.2C dataset, cluster annotation was manually curated by inspecting marker genes and SingleR (v1.8.1) classification (*60*), using as reference the dataset from Sakurai et al (*23*). As a result, we obtained a two-layer annotation, a more granular one referred to as Classification variable and a less granular one named Population to ease data interpretation. For the Fig.2H dataset, cell annotation was performed with scGATE (v1.6.2) R package (*61*) using the Fig.2C dataset as reference.

To assess the effect of expansion culture on mPB CD34+ cells we identified genes significantly deregulated across culture timepoints starting from uncultured (day0) cells to day4 and day8 expansion timepoints. To correct for differences in cell population abundances, we performed all comparisons across timepoints within each cell population (Classification label). First, we identified differentially expressed genes between timepoints (day4 vs day0; day8 vs day4 and day8 vs day0) by the FindMarker function from Seurat R package. According to fold change direction we defined different patterns of modulation that could be simplified in up or down regulated across culture (consistently up/down, early or late up/down according to day4 and day8 significance and direction). We next looked for a more comprehensive set of genes which could summarize a global culture effect across most populations, and selected sets of common up and down regulated genes shared in at least 4 populations out of 7 identified by our Population label. Lists of common up/down regulated genes across culture were used as input for ORA using clusterprofiler R package (v4.7.1) (*58*). Enrichment terms (p.adj < 0.05; Benjamini-Hochberg correction) were then plotted with the cnetplot function provided in the enrichplot r package (v1.14.2). Further details and source code are provided in the manuscript’s GitLab repository.

To evaluate the transduction efficiency of MPS patient samples, we assessed for expression of WPRE (LV specific gene) transcripts as surrogate of the IDUA LV-derived transgene, which itself could not be discriminated by IDUA transcripts originating from the native gene. Alignment and quantification of WPRE transcripts was ensured by adding the WPRE sequence to the reference human genome in cellranger (10X Genomics). Transduction efficiency (TE) was calculated as the fraction of cells bearing the WPRE transcripts within each sample and/or cell population.

### Statistical Analysis

The number of biologically independent samples, animals or experiments is indicated by n. For some experiments, different HSPC donors were pooled to account for donor-related variability and to achieve a sufficient number of cells required for the experiment. In all studies, values are expressed as means ± SEM, unless otherwise indicated.

Two-tailed tests were performed throughout the study. The Mann–Whitney test was performed to compare two independent groups, while in the presence of more than two independent groups, the Kruskal–Wallis test followed by post hoc analysis using Dunn’s test was used. In all box plots shown in the figure, the whiskers represent the minimum and maximum values of the data. Throughout the manuscript significance levels are coded as follows: *p < 0.05, **p < 0.01, ***p < 0.001.

## Data and materials availability

Lentiviral barcoding and insertion site analysis raw data were deposited on ENA under accession codes PRJEB76976, PRJEB77101, PRJEB77102 and PRJEB94025. Bulk RNA sequencing raw data were deposited at GSE292605 GEO repository. Single-cell RNA sequencing raw data were deposited under GSE292604 and GSE292606 GEO repositories for Fig. 2H and Fig. 2C datasets respectively. Code used to analyze data can be found in the following GitLab repository: [http://www.bioinfotiget.it/gitlab/custom/zonari_mpbhscexp_2025]

Materials can be made available upon signing a material transfer agreement with standard provisions, except UM171 (requests to be made to ExCellThera) and patient material/ data from the MPS1-H study (requests to be approved by OTL).

## Acknowledgements

The authors thank members from the Gentner lab (former lab in Milan and present lab in Lausanne) for help with experiments, discussion and insight; Fractal facility personnel for cell sorting; the center for Omics sciences (COSR) for advice and assistance with genomic sequencing; Cristina Tresoldi and the OSR biobank for sample collection and storage; the Pediatric and Bone Marrow Transplant Unit personnel, in particular Maria Ester Bernardo and Alessandro Aiuti, for helpful discussions and advise on the MPSI-H trial; co-investigators from the XPAND consortium, in particular Luigi Naldini, Raffaella Di Micco and Samuele Ferrari for helpful discussions. We furthermore thank Adam Wilkinson for helpful discussions on the ChemD medium. We thank Fabrizio Benedicenti for his support in designing the LV barcode amplification strategy.

## Author contributions

Conceptualization: EZ, MMN, MB, BG

Formal analysis: EZ, MMN, MB, MVo, BG

Methodology: EZ, MMN, MB, MVo, SZX, JED, BG

Investigation: EZ, MMN, MB, MVo, FV, GD, CC, IG, BM, LO, IV, MVe, LH

Resources: MVo, FT, LO, DL, IM, EM, BG

Data curation: MB

Visualization: EZ, MMN, MB, BG

Funding acquisition: EZ, BG

Project administration: EZ, MMN, MB, GD, BG

Supervision: EM, BG

Writing – original draft: EZ, MMN, MB, BG

Writing – review & editing: EZ, MMN, MB, MVo, SZX, JED, EM, BG

