## SupplementalMaterials for "Ex Vivo Expansion of Hematopoietic Stem and Progenitor Cells from Human Mobilized Peripheral Blood for Gene Therapy Applications"

### Supplemental Figure 1

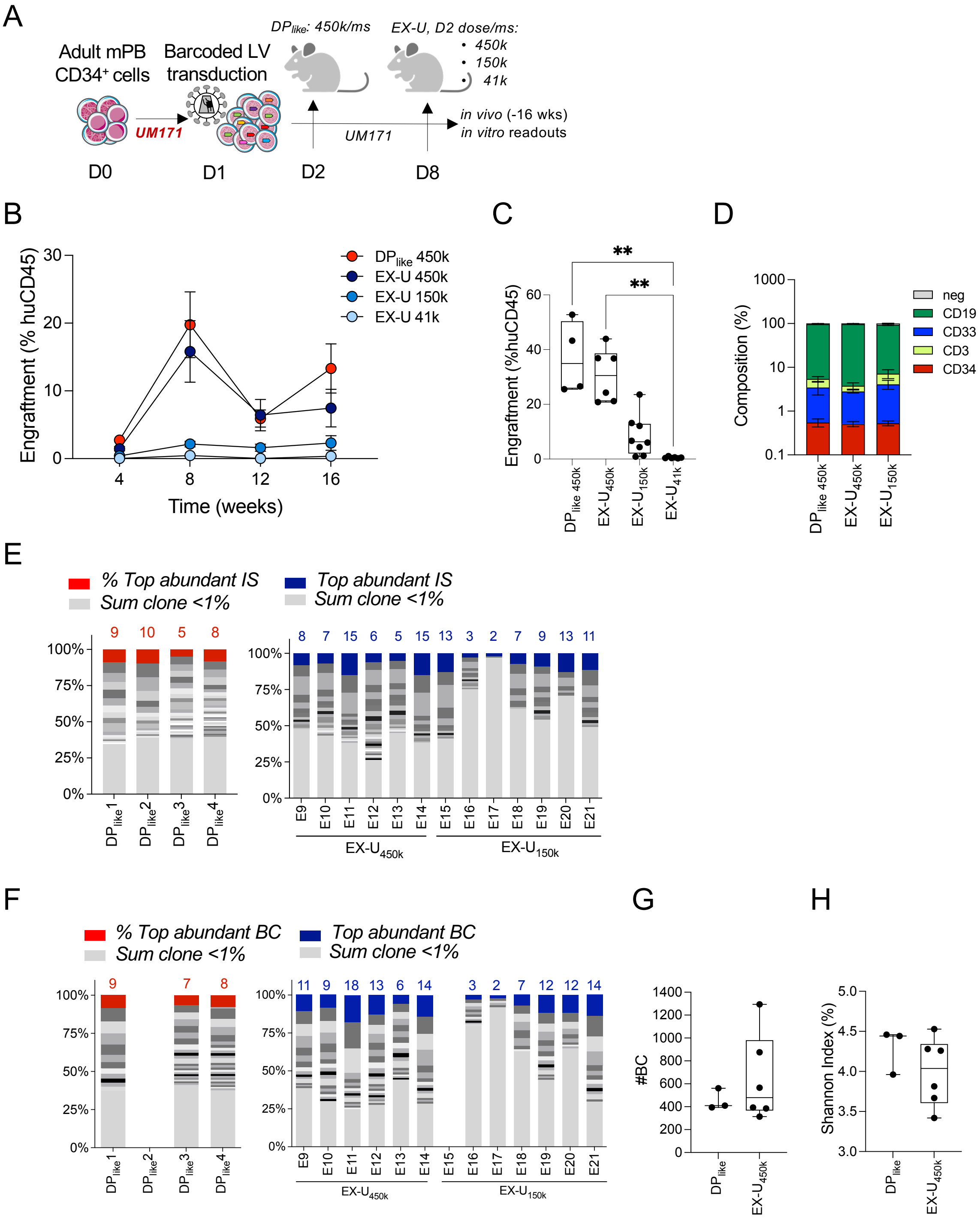

**Supplementary Fig.1 (related to Fig.1).** (**A**) Schematic representation of the experimental design. CD34+ cells from the mPB of an adult healthy donor were transduced with a purified lentiviral vector expressing a highly diverse barcode and expanded in the presence of UM171 (EX-U) or not ($DP_{like}$). The indicated doses ($k = x10^3$) of $DP_{like}$ and EX-U cells were xenotransplanted. (**B**) Percentage of human CD45+ cells (mean ± SEM) present in the peripheral blood (PB) of $DP_{like}$ (n=4), $EX\text{-}U_{450k}$ (n=6), $EX\text{-}U_{150k}$ (n=8) and $EX\text{-}U_{41k}$ (n=6) at the indicated number of weeks post-transplant. (**C**) Percentage of human CD45+ cells present in the bone marrow (BM) of $DP_{like}$ (n=4), $EX\text{-}U_{450k}$ (n=6), $EX\text{-}U_{150k}$ (n=8) and $EX\text{-}U_{41k}$ (n=6) at 16 weeks post-transplant (Kruskal-Wallis test with Dunn's multiple comparison). (**D**) Lineage composition (mean ± SEM) of the human $CD45^+$ cell graft in the BM at 16 weeks post-xenotransplantation. B cells: $CD19^+$; myeloid cells: $CD33^+$; T cells: $CD3^+$; $CD34^+$: HSPC cells; lineage negative: $CD19^-CD33^-CD3^-CD34$-. (**E**) Relative abundance clonality analysis. Each bar represents the relative abundance of insertion sites from individual mice pertaining to the $DP_{like}$ (left) or $EX\text{-}U_{450K}$ and $EX\text{-}U_{150K}$ groups (right). Bottom light grey stacks indicate the sum of insertions with <1% relative abundance, middle stacks indicate insertions with >1% abundance, where the top abundant insertions are highlighted in red or blue respectively. Number of retrieved top abundant insertions are reported above each column. (**F**) Same as in E, using the unique barcode as a clonal readout. (**G**) Box plots indicating number of unique barcodes. Each dot represents an individual mouse. (**H**) Box plots indicating the Shannon diversity index as measure of clonal population diversity analyzed by barcode. Each dot indicates an individual mouse of the indicated group.

#### Supplemental Figure 2

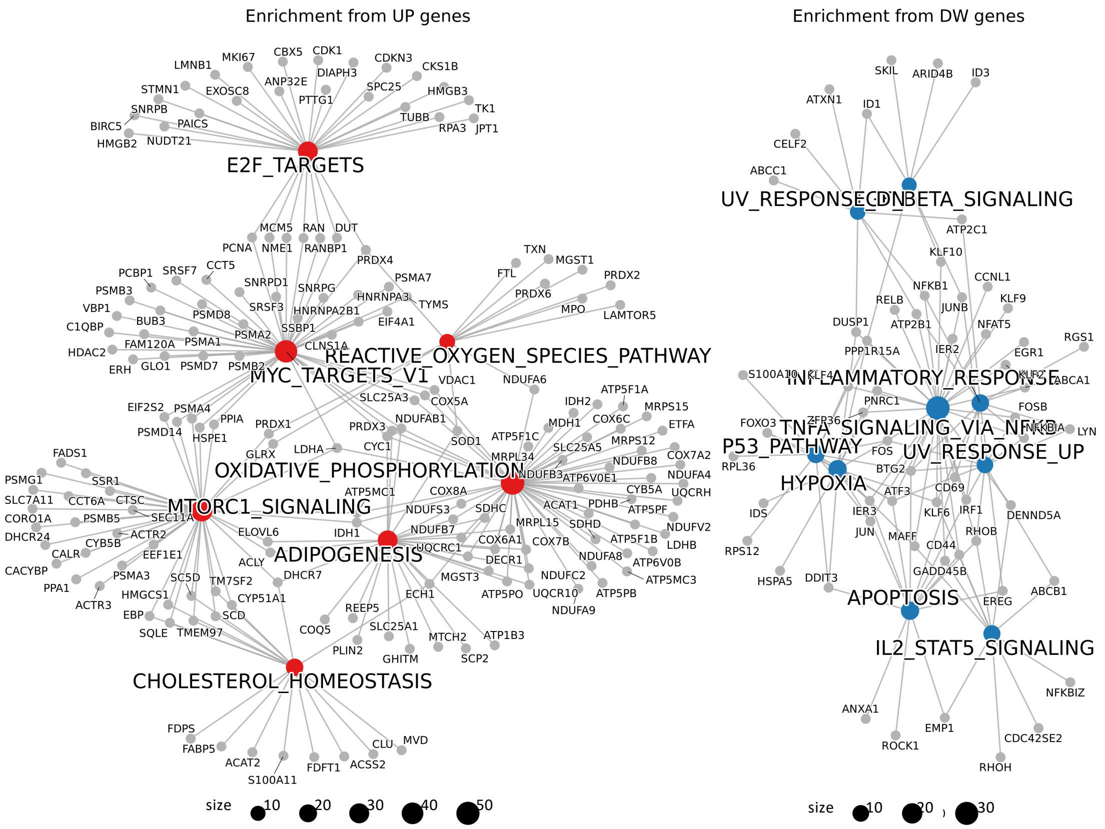


**Supplementary Fig. 2 (related to Fig.2).** ORA with MSigDB hallmark pathways enriched from commonly up- or down-regulated genes over time (from uncultured to day 4 [early] or day 8 [late]) across populations. The cnetplots display enriched terms and their associated genes: upregulated genes are shown in the left plot, and downregulated genes in the right plot. Dot size indicates the number of genes associated with each term. For further details, see the scRNA-seq Methods section.

#### Supplemental Figure 3

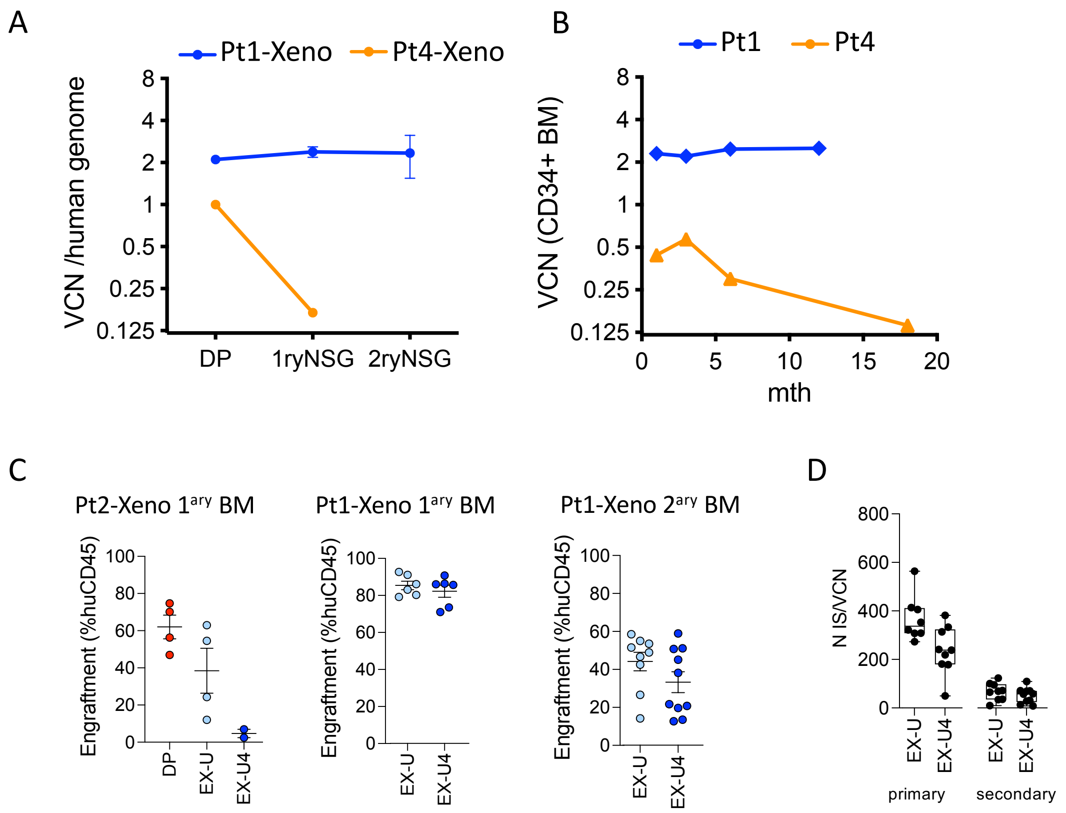


**Supplementary Fig. 3 (related to Fig.2 and Fig.4).** (**A**) Transduction levels in the drug product (DP) (as measured by vector copy number analysis) for MPS patients Pt1 and Pt4 after 14-day in vitro culture or after xenotransplantation of the DP into primary NSG mouse recipients (1ryNSG; Pt1-Xeno: n=5 mice; Pt4-Xeno: n=8 mice) and after secondary transplantation (2ryNSG; Pt1-Xeno: n=3 mice) (mean ± SEM). (**B**) VCN in BM CD34+ cells in the patients over time following gene therapy (subset of the data published in Gentner et al *(30)*, Figure S7). (**C**) Xenotransplantation of expanded DPs from MPS patients Pt1 and Pt2, expanded either in the presence of UM171 (EX-U, see Fig.1D,E) or a combination of UM171 and 4HPR (EX-U4). Human CD45+ cell engraftment is shown in the bone marrow of primary ($1^{ary}$) recipients (Pt2-Xeno, 17 weeks; Pt1-Xeno, 12 weeks. Endpoint analysis) and secondary ($2^{ary}$) recipients of the human graft harvested from the bone marrow of $1^{ary}$ recipients from Pt1-Xeno (12 weeks endpoint). (**D**) Number of unique insertion sites normalized on vector copy number found in the BM of $1^{ary}$ and $2^{ary}$ recipient mice transplanted with the expanded DP from Pt1. See also Fig.1L).

**SUPPLEMENTARY DATA FILES DESCRIPTION:**

**Data file S1:** LV Barcoding data shown in Fig S1F. It includes 7 sheets. Sheet1 describe the metadata associated to samples under analysis including group, VCN, % of engraftment and % of transduction (NGFR). This sheet is the sample sheet provided to the barcode data analysis script and includes the cutoff and MaxN variables representing the minimal % of abundance to retain a barcode and the number of max mice that can share a barcode within group to be retained in the downstream analysis. Sheet2 show the raw data produced by barseq software in tabular format. Sheet3 includes the filtered version of sheet2 according to used filters. Sheet4 shows the complete processed data provided by the data analysis pipeline with all variables included. Sheet5 shows the intragroup sharing percentage (1 mouse vs all intragroup). Sheet6 shows a summary of number of barcodes identified in each sample/group. Sheet 7 shows Shannon diversity indexes produced with vegan R package.

**Data file S2:** scRNAseq data from Fig2C dataset. Sheet1 describes some basic metrics including median values of number of UMIs and expressed genes by cells (nCount and nFeatures) as well as the % of mitochondrial genes per cell. Sheet 2 shows the cells distribution across samples and cell types. It includes the cell type % values within each sample used as input for generation of Fig.2C and Fig.2F. Sheet3 includes the cell type annotation schema according to cluster resolutions present in the deposited dataset metadata. Sheet 4 to 7 shows the markers genes according to different granularity of clustering and cell type annotations.

**Data file S3:** scRNAseq data from Fig2H dataset. Sheet1 describes some basic metrics including median values of number of UMIs and expressed genes by cells (nCount and nFeatures) as well as the % of mitochondrial genes per cell. Sheet 2 shows the cells distribution across samples and cell types. Sheet 3 and 4 shows the WPRE pos and neg cells distribution across timepoints and cell types for Pt1 and Pt4 respectively.

**Data file S4:** LV Barcoding data shown in Fig 3. It includes 8 sheets. Sheet1 describe the metadata associated to samples under analysis including group, VCN, % of engraftment and % of transduction (NGFR). This sheet is the sample sheet provided to the barcode data analysis script and includes the cutoff and MaxN variables representing the minimal % of abundance to retain a barcode and the number of max mice that can share a barcode within group to be retained in the downstream analysis. Sheet2 and Sheet3 show the raw and filtered data produced by barseq software in tabular format. Sheet4 represents the filtered version provided by the data analysis pipeline with all variables included. Sheet5 shows the intragroup sharing percentage (1 mouse vs all intragroup). Sheet6 shows a summary of number of barcodes identified in each sample/group. Sheet 7 shows the Shannon diversity indexes produced with vegan R package.

**Data file S5:** Similar structure to Data file S4. Groups in this data file are SCGM (EX-U) with 1$^{st}$ & 2$^{nd}$ transplant samples and SCGM (EX-US) with 1$^{st}$ and 2$^{nd}$ transplant samples.

**Data file S6:** Similar structure of Data file S4, but with TD timing samples from DP-like-U1 to EX-U1, EX-U2 and EX-U3 groups.

**Data file S7:** BulkRNA seq analysis 72h vs 24h. The first 2 sheets show the differentially expressed genes in the two batches (1vs1). In the 3$^{rd}$ and 4$^{th}$ sheets the concordant up and down regulated genes in the comparison 72h vs 24h across the two different batches. Sheet5 describes the enriched terms from ORA using the reactome database. Sheet 6 shows the GSEA results from integrated batches using custom senescence reference signatures. Further details are available in the methods section and in the GitLab repository to be released upon publication.
